# Physical interactions trigger *Streptomyces* to prey on yeast using natural products and lytic enzymes

**DOI:** 10.1101/2023.06.15.545052

**Authors:** Keith Yamada, Arina Koroleva, Heli Tirkkonen, Vilja Siitonen, Mitchell Laughlin, Amir Akhgari, Guillaume Mazurier, Jarmo Niemi, Mikko Metsä-Ketelä

## Abstract

Microbial predators obtain energy from killing other living cells. In this study, we present compelling evidence demonstrating that widely distributed *Streptomyces* soil bacteria, typically not considered as predators, possess the ability to detect and prey on *Saccharomyces cerevisiae*. Using fluorescence microscopy, we observed that predation is initiated by physical contact between *Streptomyces lavendulae* YAKB-15 and yeast cells. Comparative transcriptomics data indicated that the interaction triggered the production of numerous lytic enzymes to digest all major components of the yeast cell wall. The production of various glucanases, mannosidases and chitinases was confirmed by proteomics and enzymatic activity measurements. In order to destabilise the yeast cell membrane and assimilate yeast, *Streptomyces lavendulae* YAKB-15 induced production of cell-associated antifungal polyenes, namely pentamycin and filipin III, and cholesterol oxidase ChoD. In response, yeast downregulated protein synthesis and attempted to enter a quiescence-like state. We show that yeast predation is a common phenomenon in *Streptomyces*, including well-characterized strains such as *Streptomyces peucetius* ATCC 27952, where the interaction led to production of 14-hydroxyisochainin. Finally, gene inactivation studies lead us to propose a multidirectional assault model harbouring numerous redundancies that are not dependant on any single individual factor. Our results provide insights into the ecological role of *Streptomyces* and highlight the utilization of predation as a mechanism to elicit the production of bioactive natural products for drug discovery.

**Significance Statement:** Soil is a rich environment for microbes, where they compete for space and resources. *Streptomyces* bacteria are well-known for their ability to synthesize natural products, particularly antibiotics, that are used in chemical defense against competing microbes. Here we show that *Streptomyces* are, in fact, predatory bacteria. Upon encountering yeast cells, *Streptomyces* initiate the production of numerous enzymes that digest the cell wall of yeast. In addition, the interaction triggers the production of natural products that destabilize the yeast cell membrane. Collectively these actions lead to the death of yeast cells and release of cellular building blocks that *Streptomyces* can use as nutrients. The work fundamentally shifts the paradigm of how *Streptomyces* are perceived within the soil microbiome ecosystem.

## Introduction

Decades of in-depth research on soil-dwelling *Streptomyces* bacteria have proven that they are a rich source of clinically utilised antimicrobials, anticancer agents, and immunosuppressants^1^. These Gram-positive, multicellular, and non-motile bacteria have a complex life-cycle; germination of spores results in formation of branching vegetative mycelium that, upon nutrient depletion, develop into aerial hyphae and ultimately to spores. Classically, antibiotic production is associated with the onset of sporulation, with bioactive compounds being secreted to defend against competing organisms^2^. The natural products of *Streptomyces* are often associated with amensalistic killing during morphological differentiation, where one organism harms another without cost or benefit.

Soil is a relatively nutrient scarce environment where carbon is trapped in complex polysaccharides and minerals, such as iron, are limited^3^. Degradation of natural polysaccharides typically requires an array of Carbohydrate-Active enZYmes (CAZymes) with different substrate specificities to break down complex mixtures of biopolymers^4^. *Streptomyces* are intimately integrated into the global carbon cycle and produce various hydrolytic enzymes to catabolise plant biomass constituents including cellulose, hemicellulose, and lignin^5, 6^. Another abundant biopolymer is chitin, which originates from exoskeletons of insects and crustaceans, and cell walls of fungi. Efficient production of CAZymes has made *Streptomyces* a major source for industrial manufacturing in biofuel production, food and cosmetic industries, and many other fields^7^.

Despite these nutritional challenges, the soil microbiome is the most biologically diverse community in the biosphere^3^. *Streptomyces*, one of the most prevalent species of Actinobacteria that populate soils, coexist with other bacteria, fungi, and plants while competing for space and resources^8^. *Streptomyces* use a multitude of extracellular mechanisms to detect and respond to the presence of competing microbes^6^. *Streptomyces* genomes harbour a high number of diverse biosynthetic gene clusters (BGCs), which are responsible for production of secondary metabolites. However, most BGCs are silent under axenic laboratory cultures^9^ and encode cryptic metabolites that require specific environmental signal(s) for activation, either by small signalling molecules or direct physical cell-cell interactions. Activation of cryptic BGCs is a promising source of new bioactive natural products^10^ and while the medicinal value of these compounds is clear, there is growing interest in exploring the ecological role of these metabolites^11^.

Close physical contact and environmental signals between microorganisms in the soil microbiome have been shown to initiate cascades of reactions that elicit the production of secondary metabolites^11, 12^. Indeed, direct physical contact between various *Streptomyces* species and several microorganisms stimulates *Streptomyces* to produce undecylprodigiosins^13, 14^. Gamma-butyrolactones and even select antibiotics are important chemical signals for intraspecies communication and regulation of gene expression in *Streptomyces*^6^. In other cases, *Streptomyces* secondary metabolites are used for inter-kingdom communication with volatile terpenes functioning as attractants of the arthropod *Folsomia candida* and the fruit fly *Drosophila melanogaster*^15, 16^ and as a warning signal to deter the predatory nematode *Caenorhabditis elegans*^17^. Fungal interactions trigger the production of volatile organic compounds and exploration, an atypical growth mode, in many *Streptomyces* species^18^. Notably, *Streptomyces* interactions with fungal partners such as *Aspergillus nidulans* can trigger production of secondary metabolites in both parties ^12, 19^.

An extension of antagonistic microbe-microbe interactions is predation, where predatory bacteria not only kill their prey, but also consume their macromolecules as nutrients^20^. Predatory bacteria are found in different phyla and have evolved diverse strategies to kill bacteria, including epibiotic and endobiotic mechanisms, where the prey cells are lysed either from the outside or inside, respectively. The predatory behaviours of the Gram-negative soil-dwelling myxobacteria have been well characterised, including gliding motility to find prey and secretion of antibiotics and bacteriolytic enzymes to assimilate target microbes^21, 22^. Yet, discourse whether *Streptomyces* antibiotic-mediated interactions are amensalistic or predatory has been controversial^23^. *Streptomyces* are not generally appreciated as predatory bacteria, particularly because they are non-motile and due to challenges in distinguishing between non-obligatory predation and amensalism. However, recent reports have shown that *Streptomyces* isolates can grow on live cells of other bacteria^23^ and cyanobacteria^24^, when no other sources of nutrients are available. Regardless, the molecular mechanisms of the *Streptomyces* predator-prey relationships have not been elucidated in detail.

Our previous studies revealed that *Streptomyces lavendulae* YAKB-15 produces the cell-associated cholesterol oxidase ChoD strictly under co-culture conditions with whole yeast cells^25^. In this study, we noted severe perturbation of *Saccharomyces cerevisiae* cell morphology and the ultimate disappearance of yeast cells in co-cultivations with several strains of *Streptomyces*. We applied a combination of confocal fluorescence microscopy, transcriptomics and proteomics, natural product discovery, enzymatic assays, and gene knock-out studies to demonstrate that physical contact elicits the production of antifungal agents and hydrolytic exoenzymes to target the cell membrane and all major components of the yeast cell wall. We further argue that *Streptomyces* is a predator and envision that intimate physical interaction could be utilised to discover novel drug leads.

## Results

### Physical contact with *S. lavendulae* YAKB-15 induces morphological defects and disappearance of yeast cells

We were interested in examining *S. lavendulae* and *Sacc. cerevisiae* interactions during co-culture dependent production of ChoD. Microscopic evaluation revealed that yeast cells adhered to *S. lavendulae* mycelium in liquid cultures by day 1, followed by disturbances in yeast cell morphology by day 7 and ultimate disappearance of yeast cells during prolonged 20 d cultures (**Fig. 1a**). We turned to time-lapse confocal microscopy (**Supplementary Video 1**) to examine co-cultures of *S. lavendulae* YAKB-15/pS_GK_ChoD, which harbours a *choD* promoter probe plasmid for expression of green fluorescent protein (GFP), and *Sacc. cerevisiae* BY25610^26^, where red fluorescent protein (RFP) is constitutionally expressed. Still captures of a one-day movie recording revealed two yeast cell populations of approximately equivalent size at 10 min that initially duplicate at an equal rate until 300 min (**Fig. 1b**, L and R insets). *S. lavendulae* mycelium was observed growing into one of the clusters of yeast cells (**Fig. 1b**, R inset) and a significant disturbance in yeast cell growth is visible at 650 min in comparison to yeast cells that are not in physical contact with *S. lavendulae* (**Fig. 1b**, L inset). Contact with *S. lavendulae* led to a drastic decrease in the RFP signal from yeast cells and the appearance of a GFP signal, indicating activation of ChoD production, only from *S. lavendulae* in physical contact with yeast at 1000 min (**Fig. 1b**, L and R insets). The disappearance of yeast cells and the extension of the GFP signal to the entire mycelium network is evident at 1490 min (**Fig. 1b**). The changes in yeast cell morphology and loss of red fluorescence are more clearly visible at higher digital magnifications (**Supplementary Video 2**). Analysis of fluorescence signal intensities of interacting and non-interacting cell populations from the same microscopy images demonstrate significant changes with a 2.3-fold increase in GFP and a 2.3-fold decrease in RFP fluorescence in interacting populations at 1100 min (**Fig. 1c**).

**Fig. 1.**
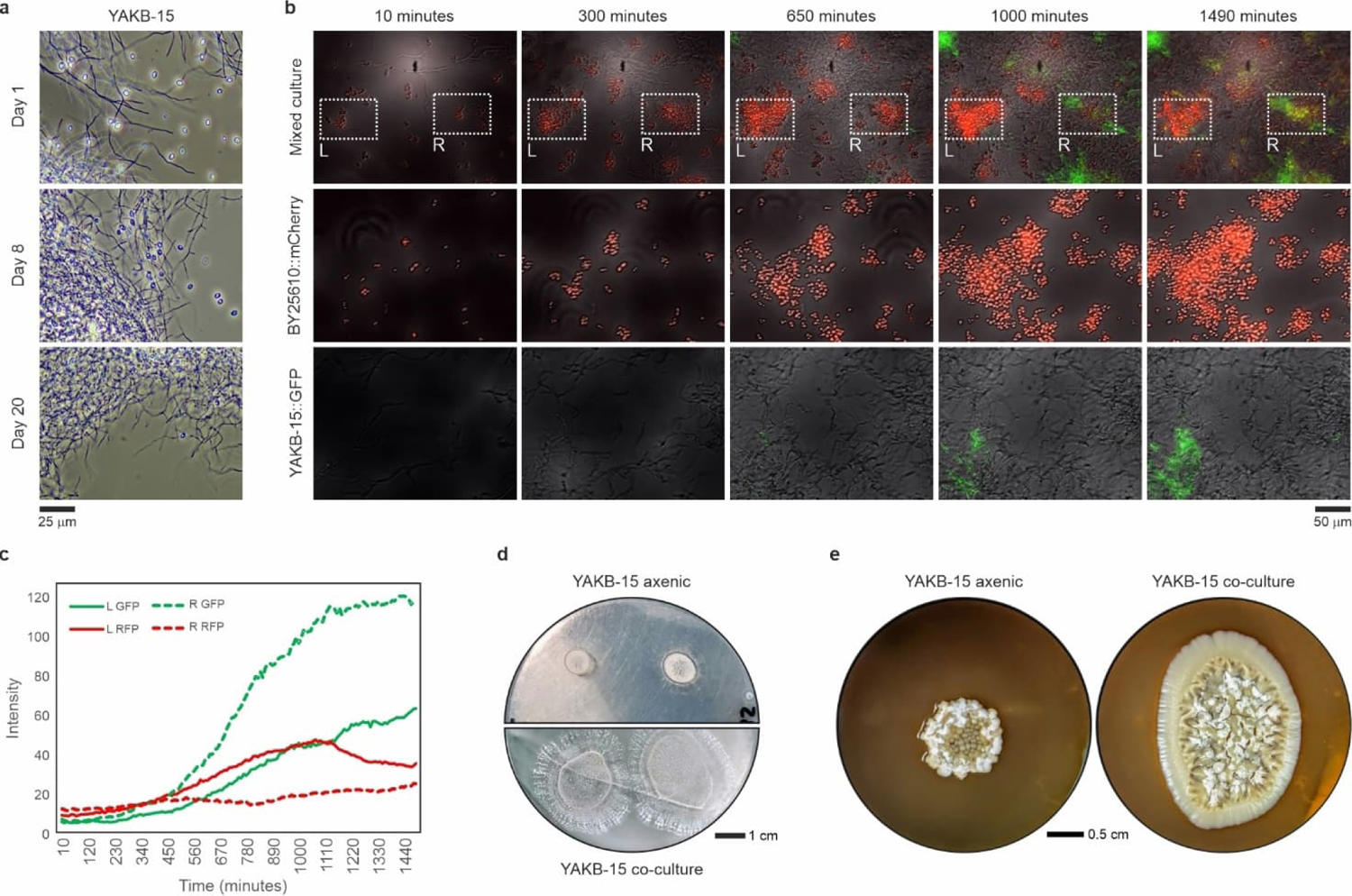
Microscopic observations of *Streptomyces*-yeast interactions. **a**, Initial observation of whole autoclaved yeast cells disappearing from a *S. lavendulae* YAKB-15 culture after 20 days. **b**, Time course confocal fluorescence microscopy of *S. lavendulae* with a *choD*-GFP promoter probe and *Sacc. cerevisiae* BY25610 constitutively expressing mCherry (RFP) from co-culture (top) and axenic cultures (middle, bottom). Two populations of yeast cells are highlighted in white boxes. The left population (L) was untouched by *S. lavendulae* and the intensity of RFP increased (**c**), while the right population (R) was interacting with *S. lavendulae* and the intensity of RFP remained low (**c**). **c**, Intensity of fluorescence measurements of the two populations over a 24-hour period. **d**,**e**, *S. lavendulae* cultured on nutrient-less and nutrient-rich agar, respectively, with and without live yeast. Notably, cultures lacked a zone of inhibition and *S. lavendulae* colonies grew larger when live yeast was present. Images are representative of triplicates.

In order to demonstrate predation, we grew *S. lavendulae* on water-agar plates, without any nutrients, both alone and on a plate with a lawn of yeast cells (**Fig. 1d**). Colony growth could be observed in axenic cultures (**Fig. 1d, top**) indicating that *S. lavendulae* can utilise agar as a carbon source, like has been reported for *S. coelicolor* A3(2)^27^, but colony size was dramatically increased under co-culture conditions (**Fig. 1d, bottom**). Equivalent results were obtained when we cultivated *S. lavendulae* on rich agar plates ideal for yeast growth (**Fig. 1e**). Inoculation of *S. lavendulae* on top of yeast cells led to *Streptomyces* consuming yeast cells. Importantly, no zone of inhibition was observed in either experiment, a characteristic of amensalism, where competitors are killed over long distances by diffusible antibiotics, (**Fig. 1d,e**), suggesting that physical contact was the mediator of predation.

### *S. lavendulae* YAKB-15 produces polyene antifungal agents upon contact with yeast

To investigate the molecular basis for the events occurring during *Streptomyces*-yeast interactions, we examined *S. lavendulae* cultures for the presence of secondary metabolites in yeast co-cultures. Comparative metabolic profiling of culture extracts revealed the appearance of two co-culture exclusive natural products, which were revealed as the known polyenes pentamycin (**1, Fig. 2**) and filipin III (**2, Fig. 2**) by 1D NMR (^1^H and ^13^C NMR) and 2D NMR (^1^H, ^1^H-COSY, HSQC, and HMBC) (Supplementary Tables 1,2 and Figs. 1-13). Interestingly, these polyenes were associated with the cell mass. Polyene antibiotics are effective antifungal agents, which mediate their mechanism of action via interactions with sterols embedded in the cell membranes that lead to destabilisation, pore formation, and cell death^28^. Of the two polyenes, **1** is used in the treatment of vaginal candidiasis, but **2** is unsuited for clinical use due to a high affinity to both fungal ergosterol and mammalian cholesterol.

**Fig. 2.**
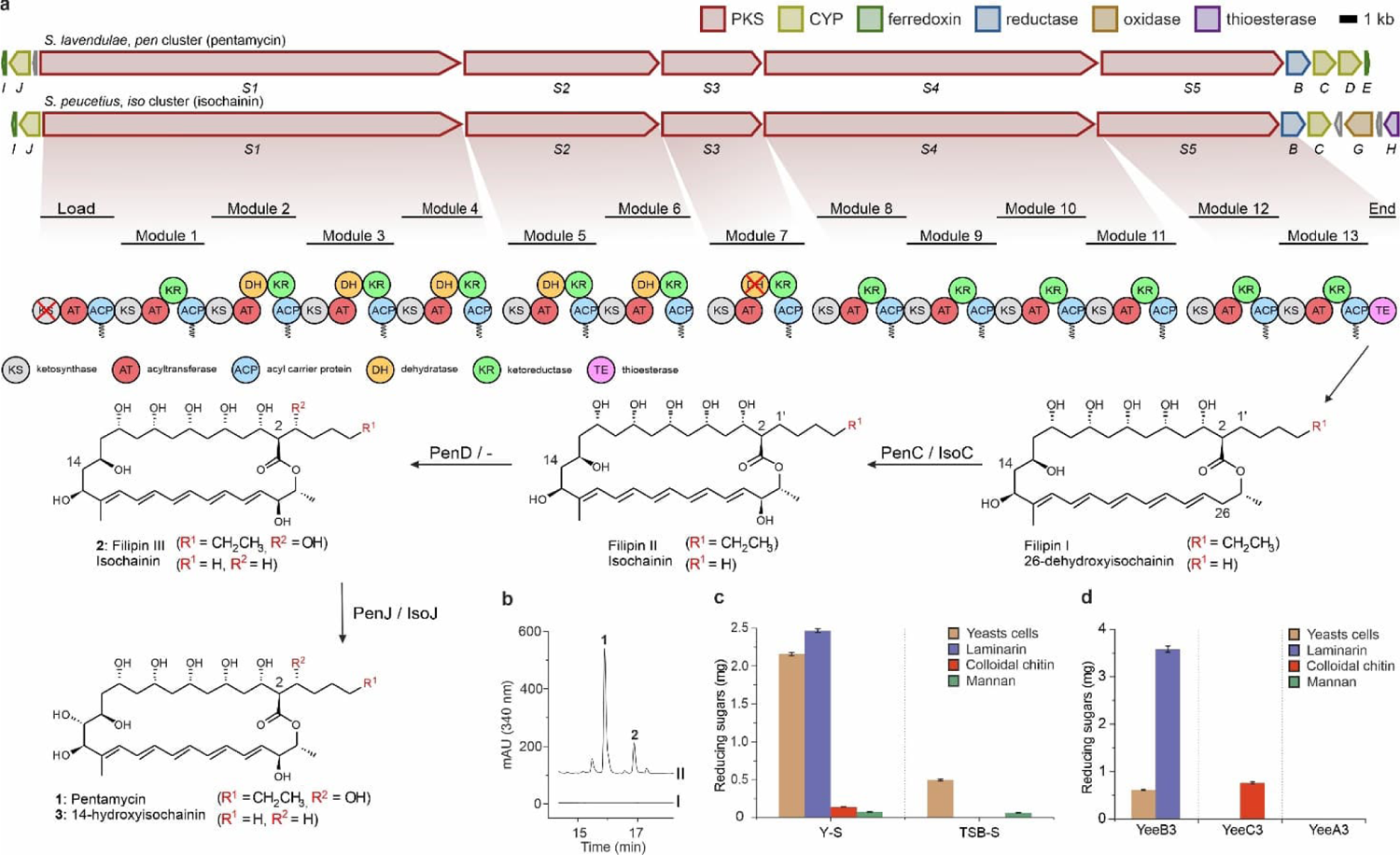
Molecular insights of the predation phenomenon. **a**, Biosynthetic gene clusters and concomitant biosynthesis of polyene antifungals (**1**: pentamycin, **2**: filipin III, **3**: 14-hydroxyisochainin) from *S. peucetius* ATCC 27952 (*iso* genes) and *S. lavendulae* YAKB-15 (*pen* genes). **b**, Chromatograms of detected polyene compounds from *S. lavendulae* cultured in different media; I: TSB medium, II: Y medium. **c**, Enzymatic activity of the *Streptomyces* secretome from cultures with whole autoclaved yeast cells (Y-S) and yeast-free (TSB-S) media, using whole autoclaved yeast cells and individual yeast cell wall components as substrates. **d**, Activities of specific yeast eating enzymes (*yee*, Fig. 3a) against whole autoclaved yeast cells and individual yeast cell wall components. Error bars indicate the standard deviation of three technical replicates.

A type I polyketide synthase BGC with domain architecture suitable for production of **1** and **2** (**Fig. 2a**) was readily identified from the genome of *S. lavendulae* due to the high (90.6 %) average nucleotide sequence identity to a pentamycin BGC from *Streptomyces* sp. S816^29^. The *pen* BGC consisted of five structural genes *penS1*-*penS5* that contained 13 type I polyketide modules with ketosynthase (KS), acyl transferase (AT), and acyl carrier protein (ACP) domains necessary for chain elongation. In addition, seven modules harboured ketoreductase (KR) domains for formation of hydroxy groups, while six modules contained a combination of KR and dehydratase domains (DH) responsible for the formation of the canonical conjugated polyene structure. Sequence analysis indicated that the DH domain in *penS3* is likely to be inactive^30^, which is also consistent with the structural analysis of pentamycin. Three genes, *penC, penD* and *penJ*, which encode cytochrome P450 mono-oxygenases, complete pentamycin biosynthesis by addition of hydroxy groups at aliphatic carbons C26, C1′^31, 32^ and C14^29^, respectively. It is noteworthy that no cluster-situated regulatory genes or putative transporters, which is in agreement with co-localization of polyenes with the cell mass, were found to reside within the BGC.

### *S. lavendulae* YAKB-15 harbours an extensive predatome of catabolic enzymes and natural products to assault yeast cells

To gain more insight into the interactions, we acquired transcriptomics data from *Streptomyces*. We proceeded to carry out RNA-Seq from axenic and co-culture samples of *S. lavendulae* with whole autoclaved yeast cells at four timepoints. Comparative transcriptomics profiling of the predatome^33^ revealed extensive changes in gene expression patterns of hundreds of CAZyme genes (**Extended Data Fig. 1**) that encode enzymes capable of lysing and digesting the polysaccharide-rich yeast cell wall^34^. We observed upregulation of different subfamilies of five α-mannosidases, four β-glucanases and seven chitinases (**Fig. 3a**). Temporal control of CAZyme gene expression correlated well with enzymatic activities required for digestion of yeast cell wall with time-course analysis indicating an initial mean 16.1-fold upregulation of secreted exo-acting α-mannosidases of the GH92 family at 12 h, followed by subsequent upregulation of intracellular GH38 family α-mannosidases at 24 h (**Fig. 3a**). The transcriptomics data revealed upregulation of the pentamycin BGC at 24 h and late-stage activation of the cholesterol oxidase *choD* expression at 48 h (**Fig. 3a**). In the biosynthetic pathway of the polyene pimaricin, the BGC embedded cholesterol oxidase *pimE* was shown to act as a signalling enzyme that triggers pimaricin production^35^.

**Fig. 3.**
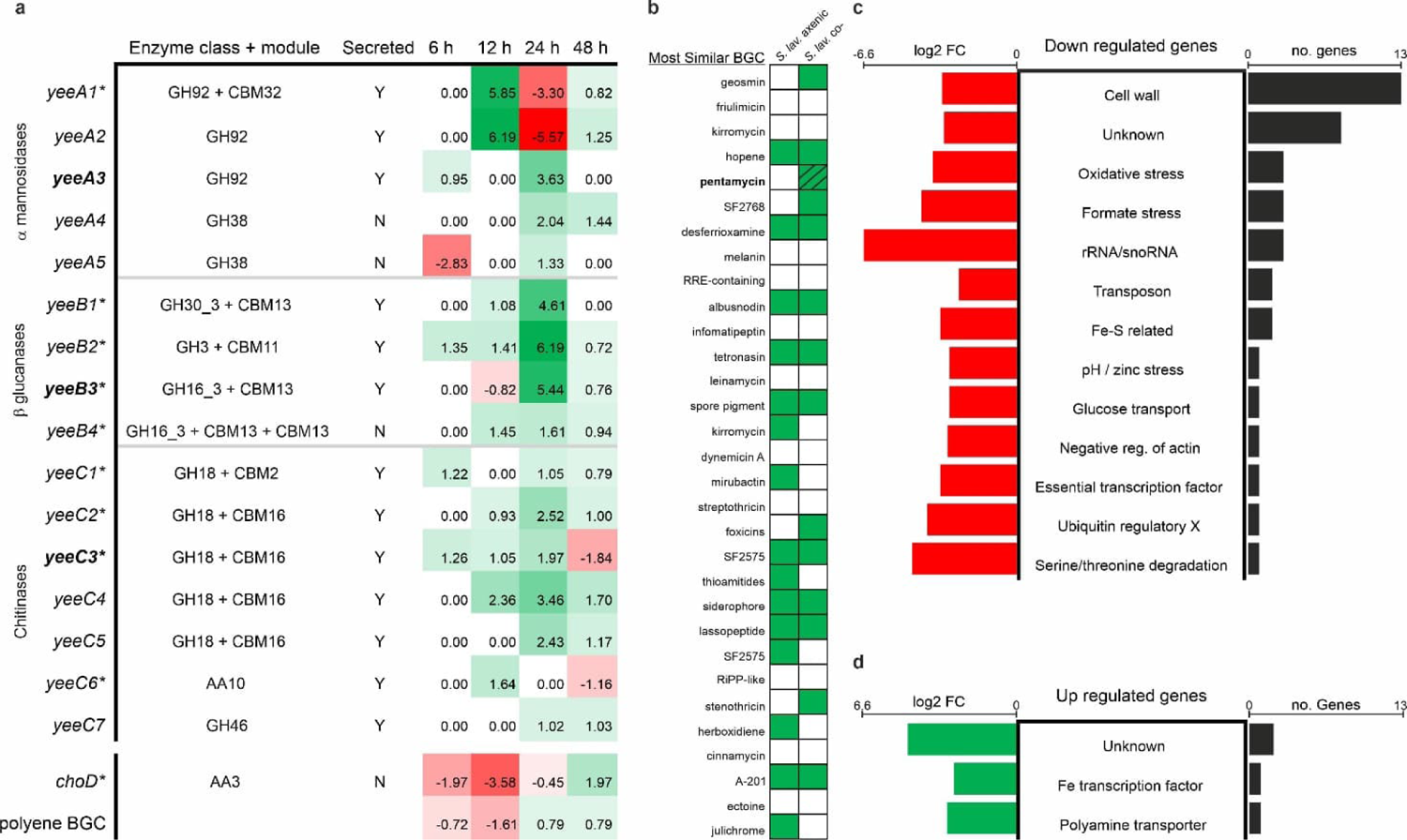
Bioinformatics, transcriptomics, and proteomics of the predation phenomenon. **a**, Comparative transcriptomics profiling reveals differential expression of enzymes targeting yeast cell wall components, as well as the *choD* and mean polyene BGC that target the yeast cell membrane, between axenic *S. lavendulae* cultures and *S. lavendulae* co-cultured with autoclaved whole yeast cells. Expression confirmed by proteomics at 24 h are marked with an *, heterologously expressed proteins are in bold (Fig. 2d), and numbers are log2 FC. **b**, Active *S. lavendulae* BGCs at 24 h (mean BGC TPM count above 90); identified compound shown with shading. **c**,**d**, Transcriptome analysis of *S. lavendulae* co-cultured with live yeast cells reveals differentially expressed yeast genes across all time points.

However, late-stage production of ChoD suggests that the function of the cholesterol oxidases may differ in these two strains and *S. lavendulae* may utilize ChoD exclusively in a catabolic role. In addition to the pentamycin BGC, transcriptomics data revealed large changes in the biosynthesis of unknown cryptic secondary metabolites (**Fig. 3b**). Five metabolic pathways were putatively activated, most notably, BGCs related to diisonitrile chalcophore SF2768-type antifungal agent, foxicin-type siderophore and stenothricin-type antibiotic, while six BGCs were putatively silenced.

We confirmed the transcriptomics data by analysis of the extracellular proteome of *S. lavendulae* by SDS-page and mass spectrometric identification of proteins. Nine out of 16 CAZymes and the cholesterol oxidase ChoD were detected from *Streptomyces*-yeast co-cultures (**Fig. 3a**). We proceeded to detect enzymatic activities against yeast cell wall components by analysing the reducing sugar content from the secretome of *S. lavendulae* cultured axenically and as a co-culture (**Fig. 2c**). Whole autoclaved yeast cells were used as a substrate to probe total hydrolytic activity, while the storage glucan laminarin, colloidal chitin, and mannan were utilised to measure specific activities. Co-cultures showed a 4.4-fold increase in total hydrolytic activity compared to axenic cultures, a high glucanase activity, and low chitinase and mannosidase activities (**Fig. 2c**). We selected the putative chitinase YeeC3 (WP_148025216.1), the glucanase YeeB3 (WP_148024776.1), and the mannosidase YeeA3 (WP_148025494.1) for heterologous protein production in *Escherichia coli* and carried out activity assays with purified proteins (**Fig. 2d**). Degradation of laminarin and chitin biopolymers into monomer components could be demonstrated with YeeB3 and YeeC3, respectively, but no activity was detected for the mannosidase YeeA3 in congruence with experiments with cell-free extracts. Collectively, our findings indicate that *S. lavendulae* harbours an extensive predatome that can degrade components of the yeast cell wall and influence cell membrane stability.

### Yeast responds by downregulating protein synthesis to enter a quiescence-like state

In contrast, transcriptome analysis of *S. lavendulae* co-cultured with live yeast indicated that yeast could not effectively defend against the attack by *S. lavendulae* and gene downregulation was the dominant response (**Fig. 3c**). The largest group of genes downregulated were related to cell wall biosynthesis, various stress responses (e.g., oxidative, formate, and pH), and strong 97-fold downregulation of ribosomal rRNA and snoRNA synthesis. The general pattern of metabolic suppression under hostile conditions suggested yeast cells have entered a quiescence-like state, reminiscent to the response of yeast to rapamycin^36^. Visually, yeast cells appear to arrest as unbudded cells with a thickened cell wall, which are characteristics of yeast quiescence^36^ (**Fig. 1a,b, Fig. 4a, Fig. 5a,b,c**). Only four genes were upregulated in yeast, where the most interesting observation was upregulation of a polyamine transporter (**Fig. 3c**). Polyamines play a key role in the yeast stress response^37^, and depletion leads to alterations of the yeast cell wall, which thickens, becomes more heterogenous and irregular in shape^38^. Membrane destabilisation by polyenes may have resulted in the loss of intracellular polyamines, which would explain the microscopic observations and changes in yeast cell morphology (**Supplementary Video 2**).

**Fig. 4.**
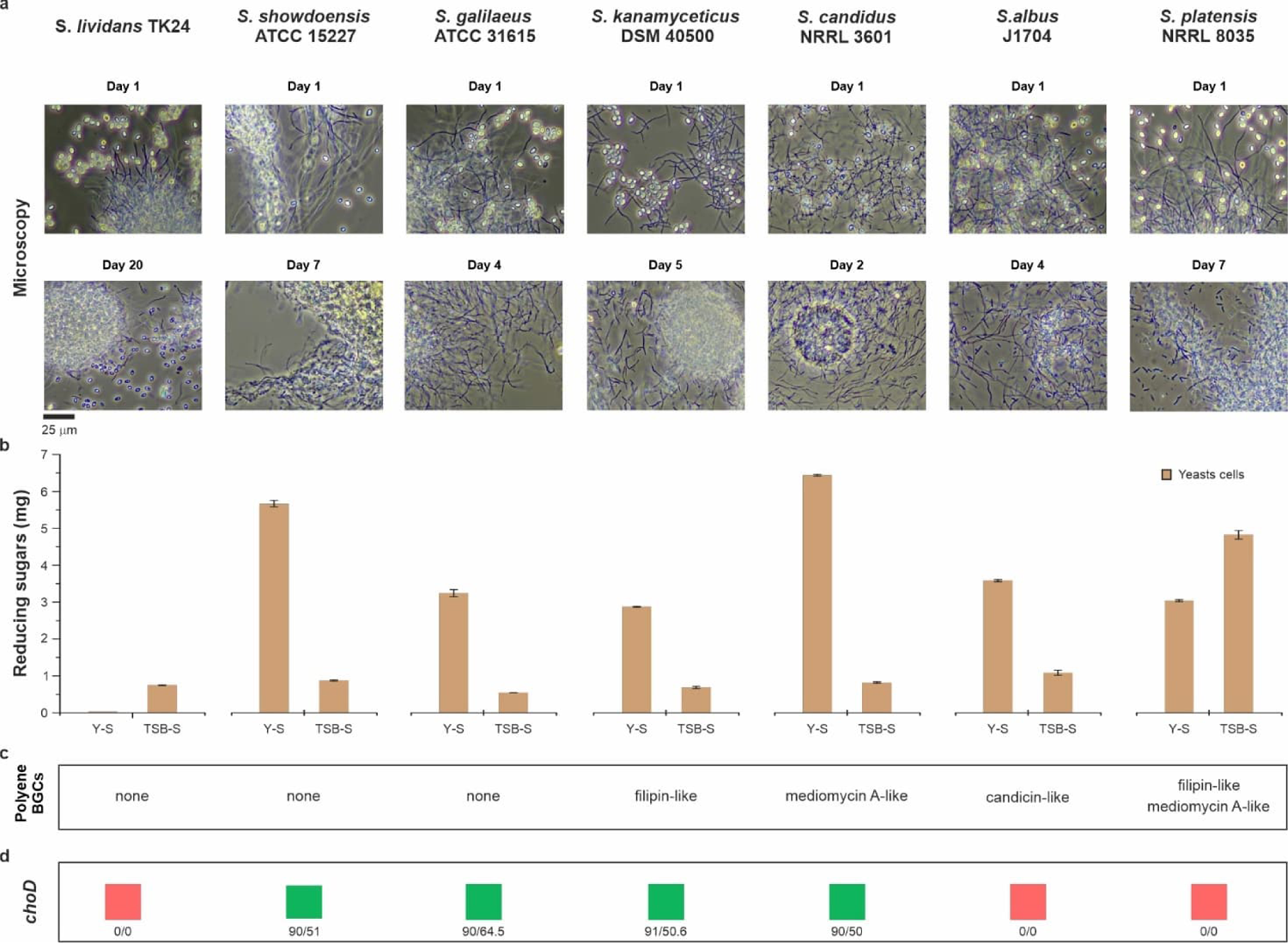
*Streptomyces* predation of yeast is widespread. **a**, Microscopic observations of seven *Streptomyces* strains during extended cultivations. **b**, Enzymatic activity against yeast cells of the *Streptomyces* secretome from cultures with whole autoclaved yeast cells (Y-S) and yeast-free (TSB-S) media, of respective strains (**a**). Error bars indicate the standard deviation of technical triplicates. **c**,**d**, Presence of polyene-type BGCs and *choD* genes in different *Streptomyces* strains (**a**), respectively. Numbers indicate coverage/identity percentages.

**Fig. 5.**
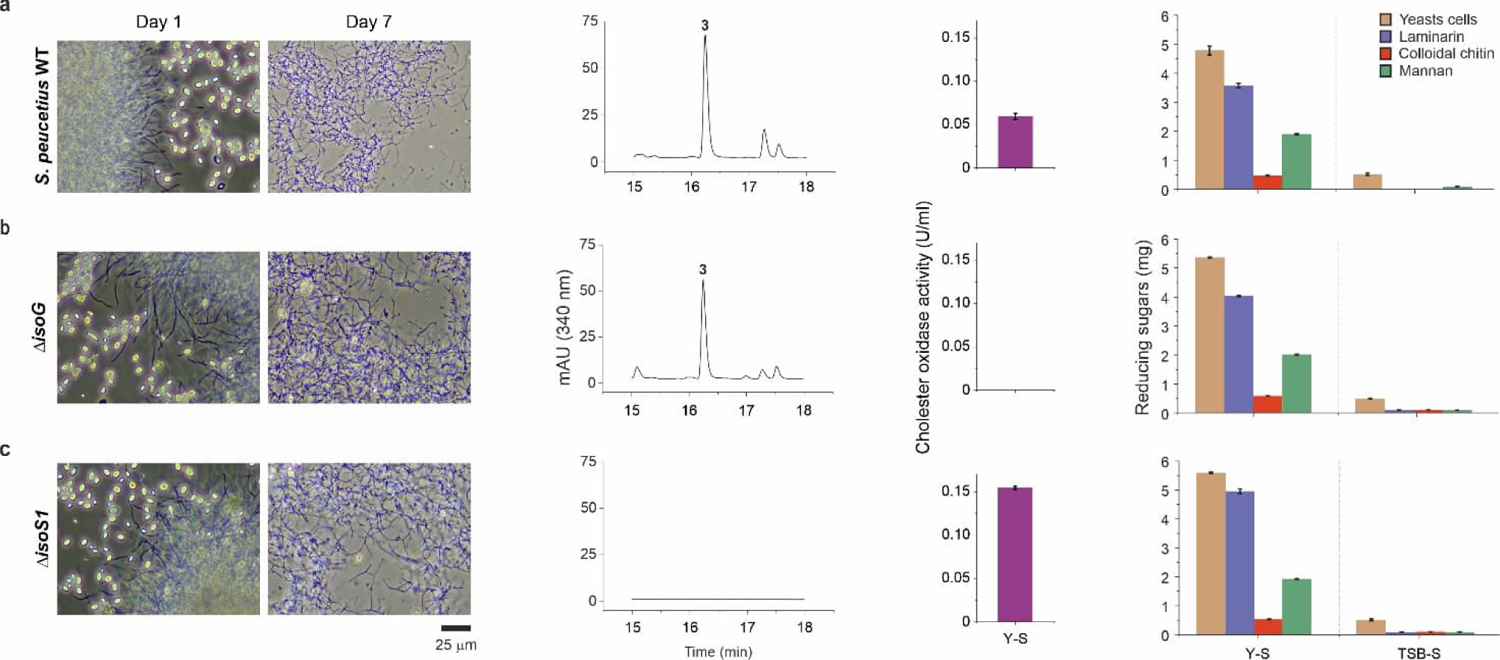
The role of the cholesterol oxidase IsoG, polyenes, and CAZymes in relation to the predation phenomenon. **a,b,c**, *S. peucetius* ATCC 27952 WT, *ΔisoG*, and *ΔisoS1*, respectively, showing (from left to right) the consumption of yeast, polyene production chromatogram, enzymatic activity of the *Streptomyces* secretome: cholesterol oxidase activity from cells grown in yeast medium (Y-S) and yeast eating enzyme activity from cultures grown in yeast (Y-S) and yeast-free (TSB-S) media. Error bars indicate the standard deviation of three technical replicates. Notably, the mutants were still able to assimilate yeast, indicating predation is not dependent on IsoG or polyenes alone.

### Genome mining reveals common yeast predatory behaviour in *Streptomyces*

We wished to interrogate if yeast predation was common in *Streptomyces* and selected eight strains with sequenced genomes for co-culture experiments (**Fig. 4a** and **Fig. 5a**). Yeast cells disappeared in seven cases and only *S. lividans* TK24 appeared to be unable to digest yeast even after 20 days (**Fig. 4a**). The kinetics of predation varied, and yeast cells were assimilated in two to seven days, depending on the strain. Hydrolytic enzyme activity from culture supernatants was notably increased in six of the strains by 3.3 to 9.3-fold under co-culture conditions, while in the case of *S. platensis* NRRL8035 the activity was reduced by 37% (**Fig. 4b**). Genome mining indicated that five of the strains harboured polyene-type BGCs, while no canonical antifungal BGCs could be clearly detected from predatory *S. showdoensis* ATCC 15127 and *S. galilaeus* ATCC 31615 strains (**Fig. 4c**). The genomic diversity was also apparent by the lack of cholesterol oxidase genes in two strains, *S. albus* J1074 and *S. platensis* NRRL 8035 (**Fig. 4d**), indicating that the mechanisms of predation may differ in different strains of *Streptomyces*. CAZyme activity was not observed in the culture supernatant of *S. lividans* TK24 and genome mining did not reveal any apparent polyene BGC or *choD* genes, which was consistent with the inability of the strain to consume yeast.

### Predatory behaviour is not dependent on a single factor in *S. peucetius* ATCC 27952

One of the strains investigated was *Streptomyces peucetius* ATCC 27952, which has been used in the manufacturing of the anticancer agent doxorubicin for nearly 50 years^39^. Analysis of co-culture extracts revealed yeast-dependent production of a metabolite (**Fig. 5a**) with the characteristic UV/Vis spectrum of polyenes with strong absorbance at 339 nm and 357 nm. Structure elucidation by 1D NMR (^1^H and ^13^C NMR) and 2D NMR (^1^H, ^1^H-COSY, HSQC, and HMBC) (Supplementary Table 3 and Figs. 14-21) revealed the compound as 14-hydroxyisochainin (**3, Fig. 2a**), which differs from **1** in a shorter fatty acid extension unit and lack of hydroxy group at C1′. The putative *iso* BGC was highly similar to the *pen* BGC with identical type I PKS domain organisation, but lacked a gene homologous to *penD*, which encodes the cytochrome P450 enzyme responsible for the instalment of the C1′-hydroxy group^31, 32^. It is noteworthy that no obvious difference could be observed in the sequence of the final extender unit AT domain that would explain differences in fatty acid-CoA selection and it may be that the choice of the starter unit is determined by intracellular concentrations similar to the biosynthesis of the lipopeptide daptomycin. No polyene transporters were detected from the BGC, but, in contrast to *S. lavendulae*, the gene cluster harboured a cholesterol oxidase *isoG* (Fig. 2a). Importantly, *S. peucetius* has not been previously described as a producer of polyenes, despite being a well-characterised strain.

Since *S. lavendulae* was not genetically tractable, we shifted our focus to *S. peucetius* to further study *Streptomyces*-yeast interactions. We showed yeast-triggered production of cholesterol oxidase, polyene, chitinase, mannosidase, and glucanase via enzymatic activities in *S. peucetius* (**Fig. 5a, Extended Data Fig. 2**). In effect, the mannosidase activity of *S. peucetius* supernatant of was nearly 10-fold higher than in *S. lavendulae* (**Fig. 2c**). To investigate whether polyenes or cholesterol oxidases are essential for *Streptomyces* predatory behaviour, we inactivated *isoG* and *isoS1* polyketide synthase gene individually in *S. peucetius* (*ΔisoG* and *ΔisoS1*). As expected, *ΔisoG* lost cholesterol oxidase activity in yeast co-cultures, and the gene inactivation did not lead to cessation of **3** production (**Fig. 5b**), which is in agreement with the delayed expression of *choD* in *S. lavendulae*. In addition, *ΔisoG* continued to produce CAZymes at levels similar to the wild-type strain. In concordance, **3** could not be detected from cultures of *ΔisoS1*, but cholesterol oxidase, chitinase, glucanase and mannosidase activities could be detected from co-cultures (**Fig. 5c**).

Surprisingly, both mutant strains were still able to assimilate yeast cells in prolonged cultures, despite their genomic deficiencies (**Fig. 5**). Together, these results suggested a redundant multipronged assault on yeast, akin to that of bacterial pathogens^40^. Bacterial pathogens, such as *Legionella pneumophila*, employ redundant virulence mechanisms with seemingly unrelated proteins and disruption of individual effector genes does not result in impaired pathogenesis.

## Discussion

Microbial predation has been known to exist for several decades and Gram-negative *δ*-*Proteobacteria* such as *Bdellovibrio*, *Bradymonas*, and *Myxococcus* have shown diverse strategies for predation^41^. Although *Streptomyces* are well known producers of bioactive antibiotics and hydrolytic exoenzymes, they have not been classically considered as predatory bacteria. *Streptomyces* were long considered to be non-motile, which precludes the wolf-pack hunting strategies enabled by the gliding motility of *Myxococcus xanthus*^22^. However, fungal interactions and volatile signalling molecules have recently been shown to trigger exploratory mobility in *Streptomyces*^18^, which occurs at similar velocities as gliding motility in *M. xanthus*^42^. The importance of mobility for predation in the soil environment can also be argued since re-wetting events can promote microbe-microbe contact^43^. Moreover, the non-motile fungus *Arthrobotrys oligospora* has been shown to trap and prey on nematodes by looped structural features of its mycelium^44^.

Here, we show that several species of *Streptomyces* can assimilate yeast cells under co-cultures. Our model suggests that physical interactions, possibly between *Streptomyces* mycelium and components of the yeast cell wall, trigger substantial changes in the transcriptome of both predator and prey. *Streptomyces* produce CAZymes capable of digesting the yeast cell wall, and cholesterol oxidase and polyenes capable of attacking the yeast cell membrane (**Fig. 6**). Similarly, the predatosome of *M. xanthus* has been shown to trigger the production of hydrolytic enzymes and natural products^45^. Moreover, our physical contact-dependent predation model provides evidence to distinguish amensalism from predation. Here, *Streptomyces* produce cell-associated cholesterol oxidase and polyenes to create spatial structuring^46^ (**Fig. 6a,b**), which limits the diffusion of the nutrients they create from killing yeast, which thus confers a direct benefit to the predator. It is noteworthy that CAZymes predicted to encode intracellular catabolic enzymes are also upregulated under co-culture conditions (**Fig. 3a**). Our multiomics and gene inactivation data suggest that neither polyenes nor ChoD are solely responsible for the predatory capabilities of *Streptomyces*, rather a redundant multipronged attack is used that is akin to bacterial pathogens^40^.

**Fig. 6.**
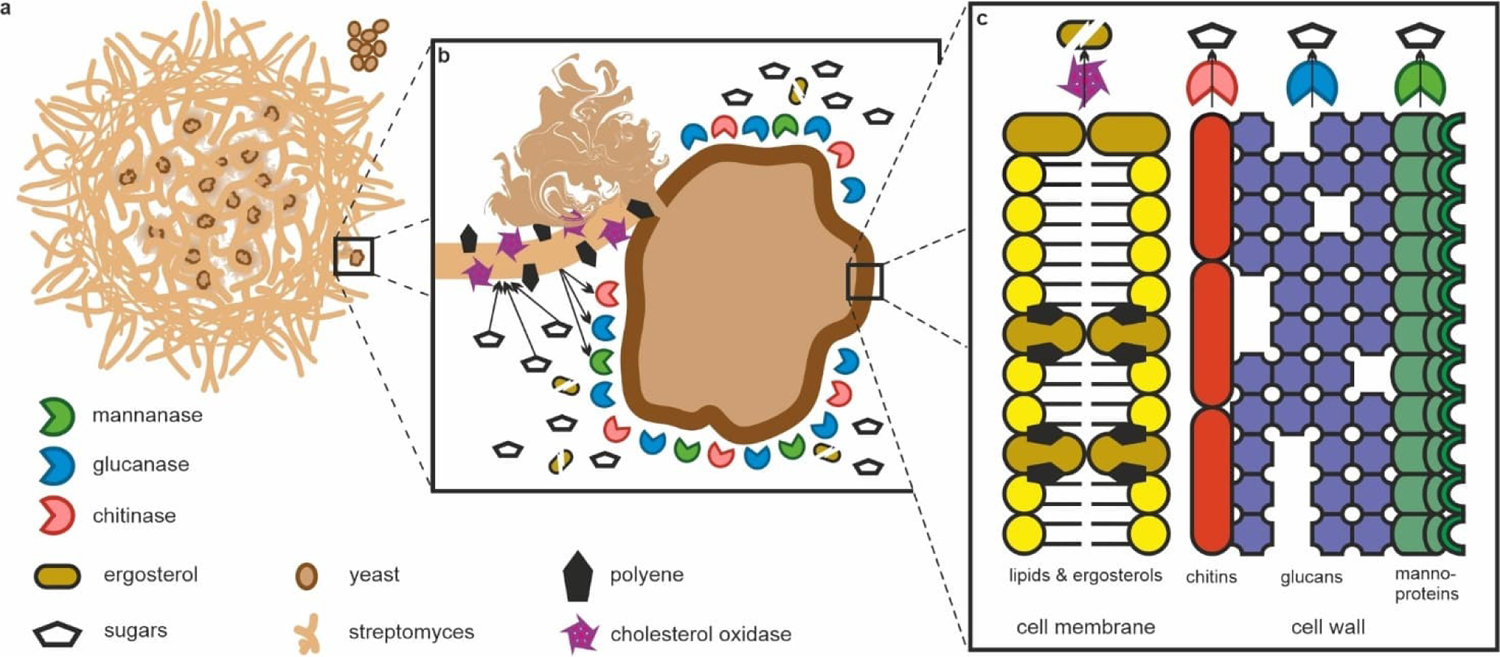
The multipronged attack model on yeast by *Streptomyces*. **a**, The attack is initiated by *Streptomyces* making physical contact with yeast, while yeast not in physical contact remain intact (Fig. 1). **b**, ChoD and the polyenes are associated with the mycelia of *Streptomyces* and are likely weapons used in the initial attack against yeast, while the CAZymes are secreted and digest the yeast cell wall (Fig. 2,3,4,5). **c**, The target of ChoD and the polyenes is the ergosterol found in the cell membrane of yeast. The target of the CAZymes are the constituents of the yeast cell wall, which *Streptomyces* is able to consume (Fig. 1).

It is noteworthy that the physical *Streptomyces*-yeast interactions triggered the production of polyenes, which are known antifungal metabolites.^28^ Our hypothesis is that physical interactions with target microbes may elicit the production of natural products specifically against the prey organism. While little is known about the molecular mechanisms of microbial predation, even in the model microbial predator *M. xanthus*, it is known that extracellular bacteriolytic enzymes and natural products play a crucial role^21, 47, 48^. Some predatory natural products even play a prey-dependent role^49, 50^. Here, we observed the activation of five BGCs, which seem to specifically activate upon contact with yeast. The upregulation of these pathways occurred during the vegetative stage of the *Streptomyces* life cycle, which is in contrast to secreted secondary metabolites associated with amensalistic killing that are typically tightly coordinated with morphological differentiation^2^. This raises the question of whether *Streptomyces* would react similarly to other microbes such as Gram-negative and Gram-positive bacteria? Co-culture dependent elicitation of cryptic BGCs for production of natural products has been widely used^9^, but the ability of *Streptomyces* to specifically detect the type of prey organism species may have been underappreciated. Our model of physical contact-dependent predation may provide a new framework for drug discovery and eliciting the production of natural products.

## Online Methods

### Microbial strains, plasmids, and culture conditions

Strains and plasmids used in this study are described in Supplementary Table 4. *Streptomyces* cultures were generally grown at 30 °C, shaking at 250-300 rpm. Strains of *Streptomyces* were cultivated in Y medium^25^ (autoclaved yeast), in SC medium^51^ (live yeast), and TSB^52^ medium as described previously. The solid media used were water agar (agar 16 g l^-1^ in reverse osmosis water), YPD^51^ and MS^52^ as previously described.

### Microscopy

Confocal fluorescence microscopy was performed using a Nikon Eclipse Ti2-E microscope with a NikonDS-Fi3 CMOS camera and a Hamamatsu sCMOS Orca Flash 4 camera attached and a Lumencor Spectra X LED light source in a temperature-controlled incubation chamber. Images were acquired using a Nikon CFI S Plan Fluor EWLD 20x/0.45 DIC N1 objective with mCherry excitation/emission bandwidths of 555 nm/632 nm and GFP excitation/emission bandwidths of 488 nm/515 nm. Images were collected using NIS-Elements AR (Nikon) and analysed using Fiji^53^.

For time-lapse imaging of *S. lavendulae*/pS_GK_ChoD^54^ (cholesterol oxidase promoter probe) and *Sacc. cerevisiae* constitutively expressing mCherry^26^, strains were first grown in SC for 48 h. The co-cultures were initiated by mixing 100 µl of *Streptomyces* with 10 µl of yeast in 1 ml of SC in a six well plate. Each strain was also grown axenically using the same respective conditions. Experiments were performed at 30 °C and images were taken every 10 min for 24 h.

Standard microscopy images were captured using a Nikon Eclipse Ci-L upright microscope with a Canon EOS RP camera attached by a TUST38C LM Direct Image C-Mount Port (Micro Tech Lab, Graz, Austria) and a DSLRCRFTC_Pro LM Digital SLR Universal Adapter (Micro Tech Lab, Graz, Austria). Solid state cultures were imaged using a WILD M3Z (Heerbrugg, Switzerland) stereomicroscope.

For plate culture imaging of *S. lavendulae* and *Sacc. cerevisiae*, strains were grown in SC for 48 h. On water agar plates, 100 µl of yeast was spread as a lawn and *Streptomyces* was spotted on top twice with 10 µl and 20 µl. *Streptomyces* was also spotted in the same way on a separate water agar plate without yeast. Water agar plates were grown at room temperature for 30 days. On YPD plates, 20 µl of yeast was spotted and allowed to dry, then 10 µl of *Streptomyces* was spotted in the middle. The same volume of *Streptomyces* was also plated on YPD without yeast. YPD plates were grown for 6 days at 30 °C.

### Chemical analysis

Compounds were extracted with methanol from the cell mass. Methanol extract was evaporated using a rotary evaporator and compounds were resuspended in H2O. Polyenes were extracted from aqueous phase with ethyl acetate, and ethyl acetate phase was dried using a rotary evaporator. Polyenes were further purified using first silica chromatography followed by semi-preparative HPLC. Fractions containing pure compounds were extracted with ethyl acetate, dried using rotary evaporator and desiccator, and resuspended in deuterated solvents (Eurisotop) for NMR measurements.

NMR spectra were recorded with 600 MHz Bruker AVANCE-III system with liquid nitrogen cooled Prodigy TCI cryoprobe or 500 MHz Bruker AVANCE-III system with liquid nitrogen cooled Prodigy BBO cryoprobe. All NMR spectra were processed in Bruker TopSpin 4.1.3 version and the signals were internally referenced to the solvent signals or tetramethylsilane. High resolution electrospray ionization mass spectra were recorded on Bruker Daltonics micrOTOF system.

HPLC-UV analyses were carried out using a SCL-10Avp/SpdM10Avp system with a diode array detector (Shimadzu) and a C18 column (2.6 μm, 100 Å, 4.6 × 100 mm Kinetex column (Phenomenex). HPLC-UV method: solvent A: 0.1 % formic acid, 15 % CH3CN, 85 % H2O; solvent B: 100 % CH3CN; flow rate: 0.5 mL/min; 0-2 min, 0 % B; 2-20 min, 0-60 % B; 20-24 min, 100 % B; 24-29 min, 0 % B. HPLC-MS analyses were carried out using an Agilent 6120 Quadrupole LCMS system linked to an Agilent Technologies 1260 infinity HPLC system using identical columns, gradients, and buffers as for HPLC-UV analyses.

Semi-preparative HPLC were carried out using a LC-20AP/CBM-20A system with a diode array detector (Shimadzu) and EVO C18, 5 μm, 100 Å, 250 x 21.2 mm Kinetex column (Phenomenex). Semi-preparative HPLC method: solvent A: 50 % 60 mM ammonium acetate – acetic acid pH 3.6, 15 % CH3CN, 35 % H2O; solvent B: CH3CN; flowrate: 20 mL/min; 0-2 min, 0 % B; 2-20 min, 0-60 % B; 20-24 min, 100 % B; 24-29 min, 0 % B. Silica chromatography was performed using high-purity grade silica (pore size 60 Å, 230-400 mesh particle size) and a gradient elution from 100:0 CHCl3/MeOH to 0:100 CHCl3/MeOH. All reagents were purchased from Sigma-Aldrich unless stated otherwise.

### Bioinformatics

The accession number of the *S. lavendulae* YAKB-15 genome is GCA_008016805.1, for the *S. peucetius* ATCC 27952 genome the accession number is GCA_002777535.1, and for the *Sacc. cerevisiae* genome the accession number is GCA_000146045.2. BGCs were identified using antiSMASH v6.1.1^55^. PKS genes were analysed using SeMPI 2.0^56^. CAZymes were functionally annotated using dbCAN2^57^. The average nucleotide identity was calculated using OrthoANIu^58^.

### Transcriptomics

Cultures for time-resolved transcriptomal profiling were performed in biological quadruplets at 30 °C and 300 rpm. For the live yeast cultures SC was used and samples were taken on days one, three, five, and seven. For the dead yeast cultures Y medium was used and samples were taken at six, 12, 24, and 48 hours. Axenic cultures were grown in the same respective medium without yeast.

RNA extraction was performed by pooling 1 ml of four independent cultures and adding 444 µl of cold STOP solution (5% phenol in ethanol). The samples were pelleted by centrifugation (5000 x *g*, 10 min, 4 °C) flash-frozen and kept at −80 °C (maximum 2 months). The cells were lysed with a mortar and pestle under liquid nitrogen and then RNA was isolated using a RNeasy Mini Kit (Qiagen) with DNase treatment. Total RNA was sent to Novogene (Cambridge, UK) for quality control (Agilent 2100), rRNA depletion (Ribo-Zero kit), library preparation (Illumina), and sequencing with NovaSeq 6000 (Illumina) to produce 2 x 150 bp reads.

All analyses were performed using the Chipster^59^ platform. The reads were manually checked using FASTQC^60^ and trimming was performed using TRIMMOMATIC^61^. The trimmed reads were aligned to the genome using BowTie2^62^ and counted using HTSeq^63^. Differential expression was performed using edgeR^64^.

Active BGCs were determined as previously described^65^, with an average transcripts per million (TPM) value of 90 as established by the detection of pentamycin.

### Proteomics

Proteomics analysis was performed on cut pieces of SDS-PAGE gel (55 kDa) from the supernatant of *S. lavendulae* cultured with (Y medium) and without (SC medium) autoclaved yeast for 1 day. In-gel digestion was performed at the Turku Proteomics Facility (Turku, Finland) according to standard protocol. The samples were analysed by LC-ESI-MS/MS using a nanoflow HPLC (Thermo Fisher) coupled to a Q Exactive HF mass spectrometer (Thermo Fisher) equipped with a nano-electrospray ionization source. Peptides were resolved on a trapping column and subsequently separated inline on a 15 cm C18 column (75 μm x 15 cm, ReproSil-Pur 3 μm 120 Å C18-AQ, Dr. Maisch HPLC GmbH, Ammerbuch-Entringen, Germany). Peptides were eluted with a 20 min gradient of 6-39%, followed by a 10 min wash stage with 100% at acetonitrile/water (80:20 (v/v)) with 0.1% formic acid.

MS data was acquired using Thermo Xcalibur v4.1 (Thermo Fisher). An information dependent acquisition method consisted of an Orbitrap MS survey scan of mass range 350–1750 *m/z* followed by HCD fragmentation for 10 most intense peptide ions. Protein identification searches were performed using Proteome Discoverer v2.5 (Thermo Fisher) connected to an in-house server running Mascot v2.7.0 (Matrix Science). Data was searched against a Swissprot *Streptomyces* database (downloaded 18.10.2021). The database search parameters were trypsin for the enzyme and 2 missed cleavages were allowed. Cysteine carbamidomethylation was set as static modification, and methionine oxidation and protein N-terminal acetylation were set as variable modifications. The peptide mass tolerance was ± 10 ppm and the fragment mass tolerance was ± 0.02 Da.

### Protein production

YeeC3, YeeB3 and YeeA3 were heterologously produced in *Escherichia coli* TOP10 strain transformed with pBADHisBΔ-yeeC3, pBADHisBΔ-yeeB3, pBADHisBΔ-yeeA3, respectively. The pre-culture was grown overnight in LB medium with 100 µg/ml of ampicillin and inoculated (1%) in 4 × 500 ml of 2xTY medium with 100 µg/ml of ampicillin. After incubation (30 °C, 4 hours, 250 rpm) to an OD600 of 0.6, cells were induced with 0.02% (w/v) ʟ-arabinose and incubated further overnight at room temperature at 180 rpm. Cultures were centrifuged (12,000 × *g*, 25 min, 4 °C) and cells pellet was resuspended in 3 volumes of wash buffer (K2HPO4 50 mM, imidazole 5 mM, NaCl 50 mM, 10% glycerol, 1% Triton X) and subsequently sonicated (Soniprep 150, MSE). Samples were centrifuged (18,000 × *g*, 30 min, 4 °C), the supernatant was collected and mixed with TALON Superflow affinity resin (GE Healthcare). After incubation for 1 hour at 4 °C, the resin was washed with wash buffer and protein was eluted with 2.5 mL elution buffer (K2HPO4 50 mM, imidazole 250 mM, NaCl 50 mM, 10% glycerol). Then the sample was buffer exchanged to the storage buffer (K2HPO4 50 mM, NaCl 50 mM, 10% glycerol) using a PD-10-column (GE Healthcare). The purified enzymes were analysed by SDS-PAGE and reducing sugars assay. *E. coli* TOP10 transformed with an empty pBADHisBΔ^66^ vector was used as a negative control for recombinant enzymes production and enzymatic assays.

### Enzymatic assays

To prepare culture supernatant for enzymatic assay, *Streptomyces* strains were cultured with autoclaved yeast (Y medium) and without (TSB medium) for 3 days. The supernatants were collected from centrifuged cultures (4000 × *g*, 10 min, 4 °C) and filtered through 0.45-μm syringe filters, followed by concentration (5-fold) in Amicon® Ultra-15 centrifugal filters (Millipore,10,000 MWCO). Concentrated supernatants were stored at 4 °C.

DNS (3,5-dinitrosalicylic acid) assay^67^ was performed to measure enzymatic activities of purified enzymes and concentrated supernatants with laminarin, colloidal chitin, mannan and 25% (w/v) yeasts cells solution as substrates. The assay was performed as follows: 50 µl of chitinase (15 μM) or glucanase (23 μM) or mannosidase (7 μM) or culture supernatant was mixed with 450 µl of the substrate (10 mg/ml in 50 mM phosphate buffer, pH 7) and incubated at 37 °C overnight. The reaction was stopped by adding 750 µl of DNS reagent and incubating samples at 95 °C for 10 min. Samples were centrifuged (12,000 × *g*, 10 min) and subjected to absorbance reading at 540 nm. Substrate sample without enzyme was used as a reaction control. Concentration (mg/ml) of reducing sugars in sample was calculated based on glucose standard curve.

ChoD assay was performed as described previously^25^ using a coupling reaction for monitoring H2O2 formation during the oxidation reaction of cholesterol. ChoD enzyme was extracted from cells with buffer (0.15% Tween 80 in 50 mM phosphate buffer solution) and was subjected to assay as follows: 120 μl Triton X-100 (0.05% in 50 mM phosphate buffer, pH 7), 10 μl ABTS (9.1 mM in MQ H2O), 2.5 μl cholesterol in ethanol (1 mg/ml), 1.5 μl horseradish peroxidase solution (150 U ml/ml) mixed with 20 μl of supernatant. ChoD assay was performed in a 96-well plate; one unit of enzyme was defined as the amount of enzyme that forms 1 μmol of H2O2 per minute at pH 7.0 and 27°C.

### Construction of *S. peucetius* mutants

The disruption of the *isoG* and *isoS1* genes was carried out via homologous recombination using the unstable multicopy pWHM3-based vector^68^, pWHM3-oriT (provided by Prof. Gilles van Wezel, Leiden University, The Netherlands). First, 1 kb of flanking regions upstream and downstream of the target genes were synthesized, followed by cloning of the apramycin resistance gene *aac(3)IV* in between flanks with *Spe*I and *Bcl*I. Then, each disruption cassette was cloned into *Hind*III and *Xba*I sites of pWHM3-oriT vector. Gene disruptions were generated by transferring the disruption construct to *E. coli* ET12567/pUZ8002 strain and conjugation with *S. peucetius* ATCC27952 strain. Double-crossover mutants were obtained after several passages of exconjugants under non-selective conditions on MS plates and further screening based on apramycin resistance and loss of thiostrepton resistance. PCR experiments were performed to confirm the deletion of the targeted genes (Supplementary Tables 5 and Fig. 22).

## Extended data (1/10)

**Extended Data Fig. 1.**
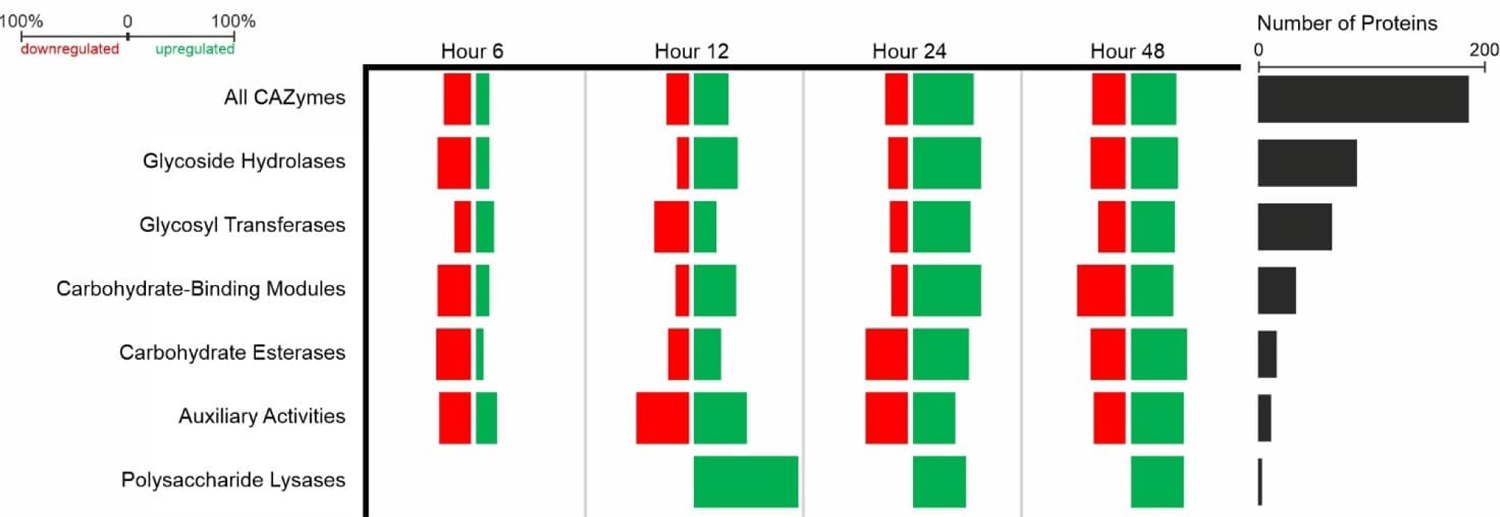
Percentage of CAZymes up or downregulated and number of CAZymes. *S. lavendulae* YAKB-15 CAZymes differentially expressed between axenic and co-cultures at several timepoint. The majority of the CAZymes are upregulated after 12 h.

**Extended Data Fig. 2.**
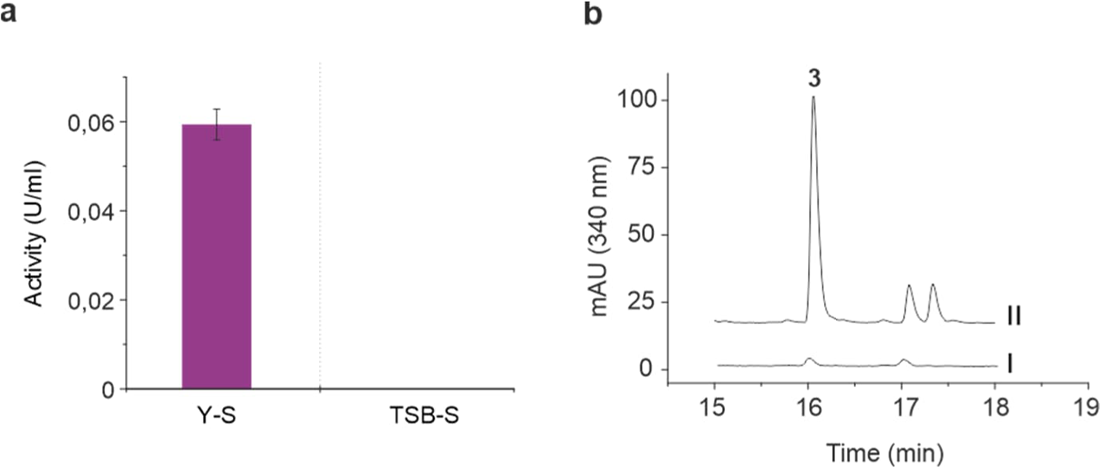
Yeast-triggered production of cholesterol oxidase and polyene in *S. peucetius* ATCC 27952. **a**, cholesterol oxidase activity from cells grown in yeast (Y*-*S) and yeast-free (TSB-S) media. Error bars indicate the standard deviation of three technical replicates. **b**, chromatograms of detected polyene (3: 14-hydroxyisochainin) from *S*. *peucetius* ATCC 27952 cultured in different media; I: TSB medium, II: Y medium.

## Data availability

The RNA-Seq data has been deposited in GEO (GSE228628). The pentamycin/filipin III BGC of *S. lavendulae* YAKB-15 and the 14-hydroxyisochainin BGC of *S. peucetius* ATCC 27952 have been deposited to the MIBiG database under the accession numbers BGC0002789 and BGC0002788, respectively.

## Acknowledgements

We thank Prof. Gilles van Wezel for providing pWHM3-oriT and Assoc. Prof. Anssi Malinen for providing *Sacc. cerevisiae* BY25610. The authors wish to acknowledge CSC – IT Center for Science, Finland, for computational resources. Mass spectrometry analyses were performed at the Turku Proteomics Facility supported by Biocenter Finland. Fluorescence imaging was performed at the Cell Imaging and Cytometry Core, Turku Bioscience Centre, Turku, Finland, with the support of Biocenter Finland. This work was funded by the Novo Nordisk Foundation Grant NNF22OC0079557 (to M.M.-K.) and the Finnish Cultural Foundation (to K.Y.).

## Author contributions

M.M-K. and J.N. conceived the project with K.Y. and A.K.; K.Y. conducted genomic, transcriptomic with A.A., and proteomic experiments and analyses; A.K. performed protein expression, enzymatic assays with A.A., and constructed mutants; K.Y., A.K., and M.L. performed the microscopy experiments; H.T., V.S., G.M. and M.L. performed the chemical analyses; M.M-K. wrote the manuscript with K.Y. and A.K.

## Competing interests

The authors declare no competing interests.

## Notes

### Competing Interest Statement

The authors have declared no competing interest.

## References

1. Atanasov, A. G., Zotchev, S. B., Dirsch, V. M. & Supuran, C. T. Natural products in drug discovery: advances and opportunities. Nat. Rev. Drug Discov. 20, 200–216 (2021).

2. Chater, K. F. Recent advances in understanding *Streptomyces*. F1000Res. 5, 2795 (2016).

3. Sokol, N. W. et al. Life and death in the soil microbiome: how ecological processes influence biogeochemistry. Nat. Rev. Microbiol. 20, 415–430 (2022).

4. Garron, M.-L. & Henrissat, B. The continuing expansion of CAZymes and their families. Curr. Opin. Chem. Biol. 53, 82–87 (2019).

5. Bentley, S. D. et al. Complete genome sequence of the model actinomycete *Streptomyces coelicolor* A3(2). Nature 417, 141–147 (2002).

6. Chater, K. F., Biró, S., Lee, K. J., Palmer, T. & Schrempf, H. The complex extracellular biology of *Streptomyces*. FEMS Microbiol. Rev. 34, 171–198 (2010).

7. Berini, F., Marinelli, F. & Binda, E. Streptomycetes: Attractive hosts for recombinant protein production. Front. Microbiol. 11, (2020).

8. Delgado-Baquerizo, M. et al. A global atlas of the dominant bacteria found in soil. Science 359, 320–325 (2018).

9. Baral, B., Akhgari, A. & Metsä-Ketelä, M. Activation of microbial secondary metabolic pathways: Avenues and challenges. Synth. Syst. Biotechnol. 3, 163–178 (2018).

10. Medema, M. H., de Rond, T. & Moore, B. S. Mining genomes to illuminate the specialized chemistry of life. Nat. Rev. Genet. 22, 553–571 (2021).

11. van Bergeijk, D. A., Terlouw, B. R., Medema, M. H. & van Wezel, G. P. Ecology and genomics of Actinobacteria: new concepts for natural product discovery. Nat. Rev. Microbiol. 18, 546–558 (2020).

12. Khalil, Z. G., Cruz-Morales, P., Licona-Cassani, C., Marcellin, E. & Capon, R. J. Inter-Kingdom beach warfare: Microbial chemical communication activates natural chemical defences. ISME J. 13, 147–158 (2019).

13. Luti, K. J. K. & Mavituna, F. Elicitation of *Streptomyces coelicolor* with dead cells of *Bacillus subtilis* and *Staphylococcus aureus* in a bioreactor increases production of undecylprodigiosin. Appl. Microbiol. Biotechnol. 90, 461–466 (2011).

14. Bikash, B. et al. Differential regulation of undecylprodigiosin biosynthesis in the yeast-scavenging *Streptomyces* strain MBK6. FEMS Microbiol. Lett. 368, fnab044 (2021).

15. Becher, P. G. et al. Developmentally regulated volatiles geosmin and 2-methylisoborneol attract a soil arthropod to *Streptomyces* bacteria promoting spore dispersal. Nat. Microbiol. 5, 821–829 (2020).

16. Ho, L. K. et al. Chemical entrapment and killing of insects by bacteria. Nat. Commun. 11, 4608 (2020).

17. Zaroubi, L. et al. The ubiquitous soil terpene geosmin acts as a warning chemical. Appl. Environ. Microbiol. 88, e00093–22 (2022).

18. Jones, S. E., et al. *Streptomyces* exploration is triggered by fungal interactions and volatile signals. eLife 6, e21738 (2017).

19. Schroeckh, V. et al. Intimate bacterial–fungal interaction triggers biosynthesis of archetypal polyketides in *Aspergillus nidulans*. Proc. Natl. Acad. Sci. 106, 14558–14563 (2009).

20. Pérez, J., Moraleda-Muñoz, A., Marcos-Torres, F. J. & Muñoz-Dorado, J. Bacterial predation: 75 years and counting! *Environ*. Microbiol. 18, 766–779 (2016).

21. Li, Z. et al. A novel outer membrane β-1,6-glucanase is deployed in the predation of fungi by myxobacteria. ISME J. 13, 2223–2235 (2019).

22. Zusman, D. R., Scott, A. E., Yang, Z. & Kirby, J. R. Chemosensory pathways, motility and development in *Myxococcus xanthus*. Nat. Rev. Microbiol. 5, 862–872 (2007).

23. Kumbhar, C., Mudliar, P., Bhatia, L., Kshirsagar, A. & Watve, M. Widespread predatory abilities in the genus *Streptomyces*. Arch. Microbiol. 196, 235–248 (2014).

24. Zeng, Y. et al. A *Streptomyces globisporus* strain kills *Microcystis aeruginosa* via cell-to-cell contact. Sci. Total Environ. 769, 144489 (2021).

25. Yamada, K. et al. Characterization and overproduction of cell-associated cholesterol oxidase ChoD from *Streptomyces lavendulae* YAKB-15. Sci. Rep. 9, 11850 (2019).

26. Breslow, D. K. et al. A comprehensive strategy enabling high-resolution functional analysis of the yeast genome. Nat. Methods 5, 711–718 (2008).

27. Merrick, M. J. A morphological and genetic mapping study of bald colony mutants of *Streptomyces coelicolor*. J. Gen. Microbiol. 96, 299–315 (1976).

28. Szomek, M. et al. Natamycin sequesters ergosterol and interferes with substrate transport by the lysine transporter Lyp1 from yeast. Biochim. Biophys. Acta BBA - Biomembr. 1864, 184012 (2022).

29. Zhou, S. et al. Pentamycin biosynthesis in philippine Streptomyces sp. S816: cytochrome P450-catalyzed installation of the C-14 hydroxyl group. ACS Chem. Biol. 14, 1305–1309 (2019).

30. Ikeda, H., Shin-ya, K. & Omura, S. Genome mining of the *Streptomyces avermitilis* genome and development of genome-minimized hosts for heterologous expression of biosynthetic gene clusters. J. Ind. Microbiol. Biotechnol. 41, 233–250 (2014).

31. Xu, L.-H. et al. Regio- and stereospecificity of filipin hydroxylation sites revealed by crystal structures of cytochrome P450 105P1 and 105D6 from *Streptomyces avermitilis*. J. Biol. Chem. 285, 16844–16853 (2010).

32. Xu, L.-H., Fushinobu, S., Ikeda, H., Wakagi, T. & Shoun, H. Crystal structures of cytochrome P450 105P1 from *Streptomyces avermitilis*: conformational flexibility and histidine ligation state. J. Bacteriol. 191, 1211–1219 (2009).

33. Pasternak, Z. et al. By their genes ye shall know them: genomic signatures of predatory bacteria. ISME J. 7, 756–769 (2013).

34. Selitrennikoff, C. P. Antifungal proteins. Appl. Environ. Microbiol. 67, 2883–2894 (2001).

35. Mendes, M. V. et al. Cholesterol oxidases act as signaling proteins for the biosynthesis of the polyene macrolide pimaricin. Chem. Biol. 14, 279–290 (2007).

36. Gray, J. V. et al. “Sleeping beauty”: Quiescence in *Saccharomyces cerevisiae*. Microbiol. Mol. Biol. Rev. 68, 187–206 (2004).

37. Valdés-Santiago, L. & Ruiz-Herrera, J. Stress and polyamine metabolism in fungi. Front. Chem. 1, (2014).

38. Miret, J. J., Solari, A. J., Barderi, P. A. & Goldemberg, S. H. Polyamines and cell wall organization in *Saccharomyces cerevisiae*. Yeast 8, 1033–1041 (1992).

39. Dhakal, D. et al. Complete genome sequence of *Streptomyces peucetius* ATCC 27952, the producer of anticancer anthracyclines and diverse secondary metabolites. J. Biotechnol. 267, 50–54 (2018).

40. O’Connor, T. J., Boyd, D., Dorer, M. S. & Isberg, R. R. Aggravating genetic interactions allow a solution to redundancy in a bacterial pathogen. Science 338, 1440–1444 (2012).

41. Mu, D.-S. et al. Bradymonabacteria, a novel bacterial predator group with versatile survival strategies in saline environments. Microbiome 8, 126 (2020).

42. Shi, W. & Zusman, D. R. The two motility systems of *Myxococcus xanthus* show different selective advantages on various surfaces. Proc. Natl. Acad. Sci. 90, 3378–3382 (1993).

43. Hartmann, M. & Six, J. Soil structure and microbiome functions in agroecosystems. Nat. Rev. Earth Environ. 4, 4–18 (2023).

44. Wang, X. et al. Bacteria can mobilize nematode-trapping fungi to kill nematodes. Nat. Commun. 5, 5776 (2014).

45. Pérez, J., Contreras-Moreno, F. J., Muñoz-Dorado, J. & Moraleda-Muñoz, A. Development versus predation: Transcriptomic changes during the lifecycle of *Myxococcus xanthus*. Front. Microbiol. 13, (2022).

46. Reintjes, G., Arnosti, C., Fuchs, B. & Amann, R. Selfish, sharing and scavenging bacteria in the Atlantic Ocean: a biogeographical study of bacterial substrate utilisation. ISME J. 13, 1119–1132 (2019).

47. Fiegna, F. & Velicer, G. J. Exploitative and hierarchical antagonism in a cooperative bacterium. PLOS Biol. 3, e370 (2005).

48. Sudo, S. & Dworkin, M. Bacteriolytic enzymes produced by *Myxococcus xanthus*. J. Bacteriol. 110, 236– 245 (1972).

49. Müller, S. et al. Identification of functions affecting predator-prey interactions between *Myxococcus xanthus* and *Bacillus subtilis*. J. Bacteriol. 198, 3335–3344 (2016).

50. Xiao, Y., Wei, X., Ebright, R. & Wall, D. Antibiotic production by myxobacteria plays a role in predation. J. Bacteriol. 193, 4626–4633 (2011).

51. Schindler, N. et al. Genetic requirements for repair of lesions caused by single genomic ribonucleotides in S phase. Nat. Commun. 14, 1227 (2023).

52. Kieser, T., Bibb, M. J., Buttner, M. J., Chater, K. F. & Hopwood, D. A. Practical streptomyces genetics. (John Innes Foundation Norwich, 2000).

53. Schindelin, J., et al. Fiji: an open-source platform for biological-image analysis. Nat. Methods 9, 676–682 (2012).

54. Akhgari, A. et al. Single cell mutant selection for metabolic engineering of actinomycetes. Metab. Eng. 73, 124–133 (2022).

55. Blin, K. et al. antiSMASH 6.0: improving cluster detection and comparison capabilities. Nucleic Acids Res. 49, W29–W35 (2021).

56. Zierep, P. F., Ceci, A. T., Dobrusin, I., Rockwell-Kollmann, S. C. & Günther, S. SeMPI 2.0—A Web server for PKS and NRPS predictions combined with metabolite screening in natural product databases. Metabolites 11, 13 (2020).

57. Zhang, H. et al. dbCAN2: a meta server for automated carbohydrate-active enzyme annotation. Nucleic Acids Res. 46, W95–W101 (2018).

58. Yoon, S.-H., Ha, S., Lim, J., Kwon, S. & Chun, J. A large-scale evaluation of algorithms to calculate average nucleotide identity. Antonie Van Leeuwenhoek 110, 1281–1286 (2017).

59. Kallio, M. A. et al. Chipster: user-friendly analysis software for microarray and other high-throughput data. BMC Genom. 12, 507 (2011).

60. Andrews S. FastQC: A quality control tool for high throughput sequence data http://www.bioinformatics.babraham.ac.uk/projects/fastqc/ (2010).

61. Bolger, A. M., Lohse, M. & Usadel, B. Trimmomatic: a flexible trimmer for Illumina sequence data. Bioinformatics 30, 2114–2120 (2014).

62. Langmead, B., Wilks, C., Antonescu, V. & Charles, R. Scaling read aligners to hundreds of threads on general-purpose processors. Bioinformatics 35, 421–432 (2019).

63. Anders, S., Pyl, P. T. & Huber, W. HTSeq—a Python framework to work with high-throughput sequencing data. Bioinformatics 31, 166–169 (2015).

64. Robinson, M. D., McCarthy, D. J. & Smyth, G. K. edgeR: a Bioconductor package for differential expression analysis of digital gene expression data. Bioinformatics 26, 139–140 (2010).

65. Amos, G. C. A. et al. Comparative transcriptomics as a guide to natural product discovery and biosynthetic gene cluster functionality. Proc. Natl. Acad. Sci. 114, E11121–E11130 (2017).

66. Kallio, P., Sultana, A., Niemi, J., Mäntsälä, P. & Schneider, G. Crystal structure of the polyketide cyclase AknH with bound substrate and product analogue: implications for catalytic mechanism and product stereoselectivity. J. Mol. Biol. 357, 210–220 (2006).

67. Miller, G. L. Use of Dinitrosalicylic acid reagent for determination of reducing sugar. Anal. Chem. 31, 426–428 (1959).

68. Vara, J., Lewandowska-Skarbek, M., Wang, Y. G., Donadio, S. & Hutchinson, C. R. Cloning of genes governing the deoxysugar portion of the erythromycin biosynthesis pathway in *Saccharopolyspora erythraea* (*Streptomyces erythreus*). J. Bacteriol. 171, 5872–5881 (1989).

