## Supplemental Information for "Physical interactions trigger *Streptomyces* to prey on yeast using natural products and lytic enzymes"

### Contents

**Table 1.** NMR data for pentamycin (**1**).
**Table 2.** NMR data for filipin III (**2**).
**Table 3.** NMR data for 14-hydroxyisochainin (**3**).
**Table 4.** Strains and plasmids used in this study.
**Table 5.** The sequence of oligonucleotides for PCR.

**Figure 1.** Comparison of solvents. $^1$H spectrum of **1** in DMSO-*d6* (blue) and MEOD (red).
**Figure 2.** $^1$H spectrum of **1** in MEOD.
**Figure 3.** $^{13}$C spectrum of **1** in MeOD.
**Figure 4.** COSY spectrum of **1** in MEOD.
**Figure 5.** HSQCDE spectrum of **1** in MeOD.
**Figure 6.** HMBC spectrum of **1** in MEOD.
**Figure 7.** HR-MS spectrum of **1**. [H-M-H]$^-$ calc. 651.3783 obs. 651.3781.
**Figure 8.** $^1$H spectrum of **2** in MEOD.
**Figure 9.** $^{13}$C spectrum of **2** in MeOD.
**Figure 10.** COSY spectrum of **2** in MeOD.
**Figure 11.** HSQCDE spectrum of **2** in MEOD.
**Figure 12.** HMBC spectrum of **2** in MeOD.
**Figure 13.** HR-MS spectrum of **2**. [H-M-H]$^-$ calc. 635.2958, obs. 635.3808.
**Figure 14.** $^1$H NMR spectrum ($CD_3OD$, 600 MHz) of **3**.
**Figure 15.** $^{13}$C NMR spectrum ($CD_3OD$, 125 MHz) of **3**.
**Figure 16.** $^1$H,$^1$H-COSY spectrum ($CD_3OD$, 600 MHz) of **3**.
**Figure 17.** $^1$H,$^1$H-COSY (**▬**) and selected HMBC (→) correlations of **3**.
**Figure 18.** HSQC spectrum ($CD_3OD$, 600 MHz) of **3**.
**Figure 19.** HMBC spectrum ($CD_3OD$, 600 MHz) of **3**.
**Figure 20.** HPLC-MS analysis of **3**.
**Figure 21.** HR-MS spectrum of **3**. [M-$H_2O$-H]$^-$ calc. 607.3482, obs. 607.3500.
**Figure 22.** Conformation of gene knockout by PCR analysis.

**Table 1.** NMR data for pentamycin (**1**).

| | **δ ppm** | **δ ppm, *J* Hz** |
|---|---|---|
| **Position** | **$^{13}$C** | **$^{1}$H** |
| 1 | 173.0 | |
| 2 | 60.5 | 2.56 dd 7.2, 9.2 |
| 3 | 73.3 | 4.18 m |
| 4 | 41.3 | 1.51 m |
| 5 | 74.2 (16) | 4.01 m |
| 6 | 45.2 | 1.48 m |
| | | 1.38 m |
| 7 | 74.0 | 4.01 m |
| 8 | 45.4 | 1.48 m |
| | | 1.38 m |
| 9 | 74.2 (23) | 4.01 m |
| 10 | 44.4 | 1.5 m |
| 11 | 71.5 | 3.97 m |
| 12 | 39.5 | 1.54 m |
| | | 1.76 ddd 3.5, 10.8, 14.0** |
| 13 | 70.4 | 3.27 ddd 1.5, 1.9, 10.8 |
| 14 | 78.3 | 3.71 dd 1.9, 9.0 |
| 15 | 80.5 | 3.89 d 9.0 |
| 16 | 138.6 | |
| 17 | 129.9 | 6.06 dd 1.1, 11.0 |
| 18 | 129.1 | 6.47 m |
| 19 | 135.5 | 6.35 m |
| 20 | 134.2 (17) | 6.36 m |
| 21 | 134.9 | 6.36 m |
| 22 | 133.7 | 6.36 m |
| 23 | 134.2 (24) | 6.31 m |
| 24 | 132.0 | 6.45 m |
| 25 | 134.4 | 6.03 dd 4.8, 14.1 |
| 26 | 73.2 | 4.10 ddd 1.1, 5.5, 6.9 |
| 27 | 75.3 | 4.84 m* |
| 28 | 18.0 | 1.29 d 6.4 3H |
| 29 | 11.7 | 1.79 d 1.0 3H** |
| 1' | 72.5 | 3.85 ddd 2.1, 8.9, 9.1 |
| 2' | 36.3 | 1.51 m |
| | | 1.38 m |
| 3' | 26.1 | 1.53 m |
| | | 1.38 m |
| 4' | 33.0 | 1.32 m |
| 5' | 23.8 | 1.33 m |
| 6' | 14.5 | 0.9 t 7.0 3H |

* Overlapping with solvent peak, ** Partly overlapping H12 and H29

**Table 2.** NMR data for filipin III (**2**).

| | **δ ppm** | **δ ppm, *J* Hz** |
|---|---|---|
| **Position** | **$^{13}$C** | **$^{1}$H** |
| 1 | 173.1 | |
| 2 | 60.5 | 2.55 dd 7.4, 9.2 |
| 3 | 73.3 | 4.18 m |
| 4 | 41.6 | 1.51 m |
| 5 | 73.8 | 3.99 m |
| 6 | 45.3 | 1.47 m |
| | | 1.37 m |
| 7 | 73.7 | 3.99 m |
| 8 | 45.4 (37) | 1.47 m |
| | | 1.37 m |
| 9 | 74.1 | 3.99 m |
| 10 | 44.3 | 1.52 m |
| | | 1.33 m |
| 11 | 71.2 | 3.98 m |
| 12 | 45.4 (41) | 1.73 m** |
| 13 | 67.6 | 3.23 dddd 1.5, 2.6, 10.9, 11.3 |
| 14 | 43.0 | 1.90 ddd 3.1, 10.7, 13.1 |
| | | 1.73 m** |
| 15 | 76.0 | 4.13 dd 10.7, 4.5 |
| 16 | 140.4 | |
| 17 | 128.5 | 6.04 dd 11.1, 1.2 |
| 18 | 129.4 | 6.49 dd 11.3, 14.1 |
| 19 | 135.1 | 6.35 m |
| 20 | 134.3 | 6.35 m |
| 21 | 134.8 | 6.35 m |
| 22 | 133.9 | 6.35 m |
| 23 | 134.4 | 6.35 m |
| 24 | 132.3 | 6.43 m |
| 25 | 134.5 | 5.98 dd 5.3, 15.0 |
| 26 | 73.75 | 4.07 ddd 1.1, 5.6, 7.4 |
| 27 | 75.2 | 4.83 m* |
| 28 | 18.1 | 1.30 d 6.3 3H |
| 29 | 11.1 | 1.77 d 0.8 3H |
| 1' | 72.6 | 3.85 ddd 2.1, 8.6, 9.3 |
| 2' | 36.3 | 1.49 m |
| | | 1.39 m |
| 3' | 26.2 | 1.52 m |
| | | 1.37 m |
| 4' | 33.1 | 1.33 m |
| | | 1.29 m |
| 5' | 23.8 | 1.33 m |
| 6' | 14.6 | 0.92 t 7.0 3H |

* Overlapping with solvent peak, ** Partly overlapping H12 and H29

**Table 3.** NMR data for 14-hydroxyisochainin (**3**).

| | **δ ppm** | **δ ppm, *J* Hz** |
|---|---|---|
| **Position** | **$^{13}C$** | **$^{1}H$** |
| 1 | 175.4 | |
| 2 | 54.4 | 2.33 ddd 3.8, 7.9, 10.6 |
| 3 | 73.6 (58) | 4.00 m |
| 4 | 42.4 | 1.41 m |
| | | 1.40 m |
| 5 | 74.1 | 4.00 m |
| 6 | 44.9 | 1.46 m |
| | | 1.36 m |
| 7 | 73.6 (61) | 4.00 m |
| 8 | 45.2 | 1.46 m |
| | | 1.36 m |
| 9 | 73.7 | 3.80 m |
| 10 | 44.2 | 1.49 m |
| | | 1.33 m |
| 11 | 71.4 | 3.97 m |
| 12 | 39.5 | 1.75 m |
| | | 1.54 m |
| 13 | 70.3 | 3.29 m* |
| 14 | 78.2 | 3.71 dd 1.8, 9.0 |
| 15 | 80.4 | 3.90 d 9.0 |
| 16 | 138.8 | |
| 17 | 129.8 | 6.06 dd 1.1, 11.3 |
| 18 | 129.3 | 6.49 dd 11.3, 14.0 |
| 19 | 135.2 | 6.33 m |
| 20 | 134.4 (35) | 6.36 m |
| 21 | 134.5 | 6.36 m |
| 22 | 133.8 | 6.33 m |
| 23 | 134.0 | 6.33 m |
| 24 | 132.0 | 6.40 m |
| 25 | 134.4 (41) | 5.98 dd 5.4, 15.3 |
| 26 | 73.5 | 4.03 m |
| 27 | 74.6 | 4.87 m* |
| 28 | 18.3 | 1.31 d 6.5 3H |
| 29 | 11.8 | 1.79 d 1.1 3H |
| 1' | 30.2 | 1.71 m |
| | | 1.57 m |
| 2' | 30.6 | 1.29 m |
| 3' | 23.7 | 1.32 m |
| 4' | 14.3 | 0.90 t 7.0 3H |

* Overlapping with solvent peak

**Table 4.** Strains and plasmids used in this study

| **Strains** | **Genotype/comments** | **Source/reference** |
|---|---|---|
| ***S. lavendulae*** | | |
| YAKB-15 | Wild-type | [1] |
| ***S. peucetius*** | | |
| ATCC 27952 | Mutant from ATCC 29050 | [2] |
| Δ*isoS1* | Δ*isoS1::apr* | This work |
| Δ*spchoD* | Δ*spchoD::apr* | This work |
| ***S. galilaeus*** | | |
| ATCC 31615 | Wild-type | [3] |
| ***S. albus*** | | |
| J1074 | Wild-type | [4] |
| ***S. showdoensis*** | | |
| ATCC 15227 | Wild-type | [5] |
| ***S. lividans*** | | |
| TK24 | Wild-type | [4] |
| ***S. kanamyceticus*** | | |
| DSM 40500 | Wild-type | [6] |
| ***S. candidus*** | | |
| NRRL 3601 | Wild-type | [7] |
| ***S. platensis*** | | |
| NRRL 8035 | Wild-type | [8] |
| ***Saccharomyces cerevisiae*** | | |
| BY25610 | BY4741ho::Nat-TEFIIpr-mCherry-ADH1ter | Dr. Anssi Malinen |
| ***E. coli*** | | |
| TOP10 | F-mcrA Δ(mrr-hsdRMS-mcrBC) Φ80lacZΔM15 ΔlacX74 recA1 araD139 Δ(araleu)7697 galU galK rpsL (StrR) endA1 nupG | Invitrogen |
| ET12567/pUZ8002 | *dam-13*::Tn9 *dcm-6 hsdM* pUZ8002 | [4] |

**Plasmids**

| | | |
|---|---|---|
| pBADΔHis | N terminal tail replaced with AHHHHHHHR | [9] |
| pWHM3-oriT | Derivative of pWHM3 | Prof. Gilles van Wezel |

**Table 5.** The sequence of oligonucleotides for PCR

| Gene | Forward | Reverse |
|---|---|---|
| *spchoD*<br>Δ*isoS1* | CAGATCGGACAAGGACAAC | GAATGACCACTGCTGTGAG |
| *spchoD*<br>*S. peucetius* ATCC 27952 | CAGATCGGACAAGGACAAC | CCAACGGCAACATCATGAC |
| *isoS1*<br>Δ*isoS1* | CAGAGGAAGAAGGGAATGGG | TAGCACGATCAACGGCAC |
| *isoS1*<br>*S. peucetius* ATCC 27952 | CAGAGGAAGAAGGGAATGGG | ATGTTGAGGTTCACGCCG |

**References for Table 4**


1. Yamada, K. *et al.* Characterization and overproduction of cell-associated cholesterol oxidase ChoD from *Streptomyces lavendulae* YAKB-15. *Sci. Rep.* **9**, 11850 (2019).
2. Dhakal, D. *et al.* Complete genome sequence of *Streptomyces peucetius* ATCC 27952, the producer of anticancer anthracyclines and diverse secondary metabolites. *J. Biotechnol.* **267**, 50–54 (2018).
3. Niemi, J. *et al.* Hybrid anthracycline antibiotics: production of new anthracyclines by cloned genes from *Streptomyces purpurascens* in *Streptomyces galilaeus*. *Microbiology* **140**, 1351–1358 (1994).
4. Kieser, T., Bibb, M. J., Buttner, M. J., Chater, K. F. & Hopwood, D. A. *Practical streptomyces genetics*. (John Innes Foundation Norwich, 2000).
5. Palmu, K. *et al.* Discovery of the showdomycin gene cluster from *Streptomyces showdoensis* ATCC 15227 yields insight into the biosynthetic logic of C-nucleoside antibiotics. *ACS Chem. Biol.* **12**, 1472–1477 (2017).
6. Zhang, S. *et al.* Establishment of a highly efficient conjugation protocol for *Streptomyces kanamyceticus* ATCC12853. *MicrobiologyOpen* **8**, (2019).
7. Zhao, G. *et al.* The biosynthetic gene cluster of pyrazomycin—A C-nucleoside antibiotic with a rare pyrazole moiety. *ChemBioChem* **21**, 644–649 (2020).
8. Osada, H., Koshino, H., Kudo, T., Onose, R. & Isono, K. A new inhibitor of protein kinase C, RK-1409 (7-oxostaurosporine). I. Taxonomy and biological activity. *J. Antibiot. (Tokyo)* **45**, 189–194 (1992).
9. Kallio, P., Sultana, A., Niemi, J., Mäntsälä, P. & Schneider, G. Crystal structure of the polyketide cyclase AknH with bound substrate and product analogue: implications for catalytic mechanism and product stereoselectivity. *J. Mol. Biol.* **357**, 210–220 (2006).

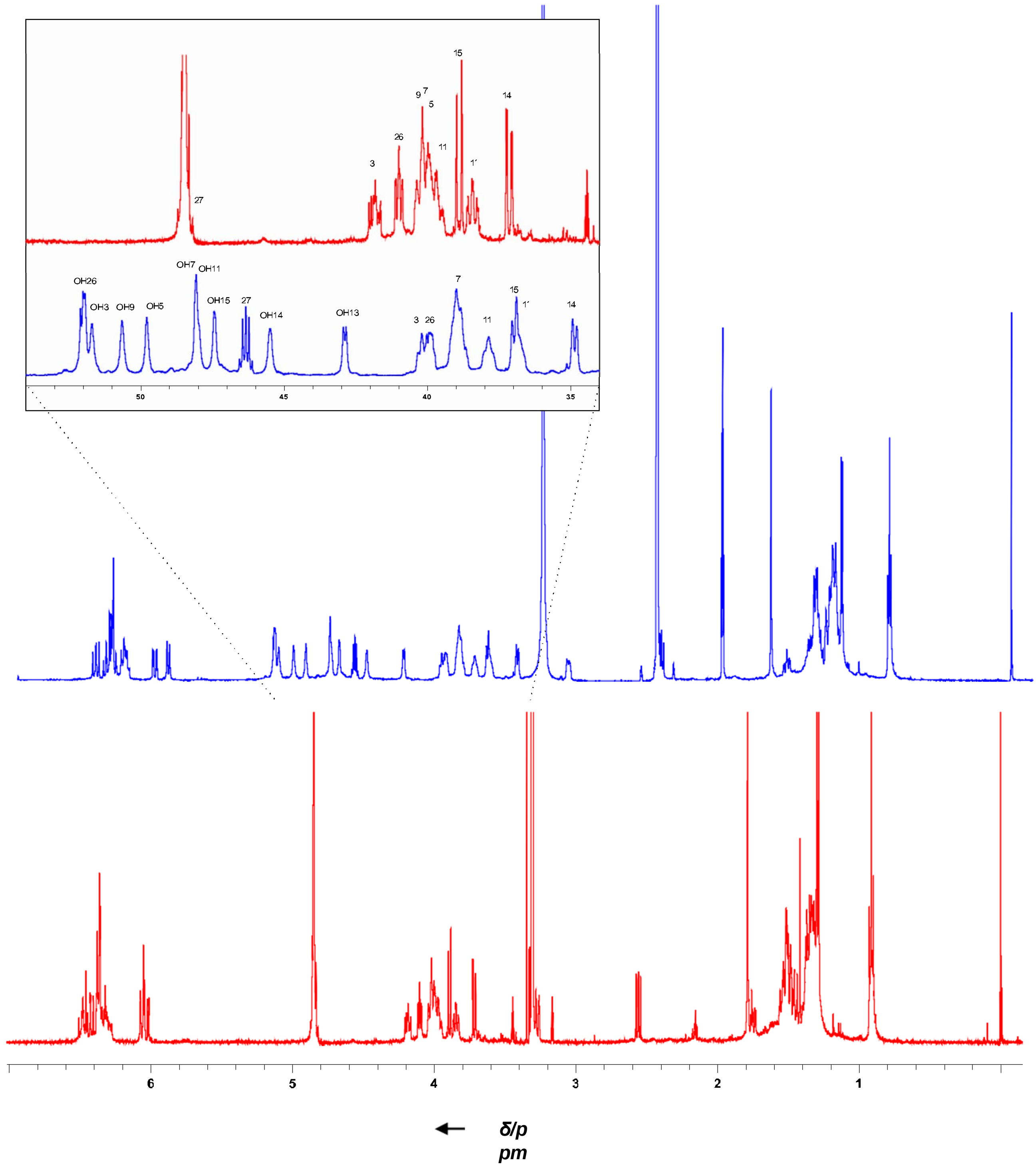


**Figure 1.** Comparison of solvents. [1]H spectrum of **1** in DMSO-*d6* (blue) and MEOD (red).

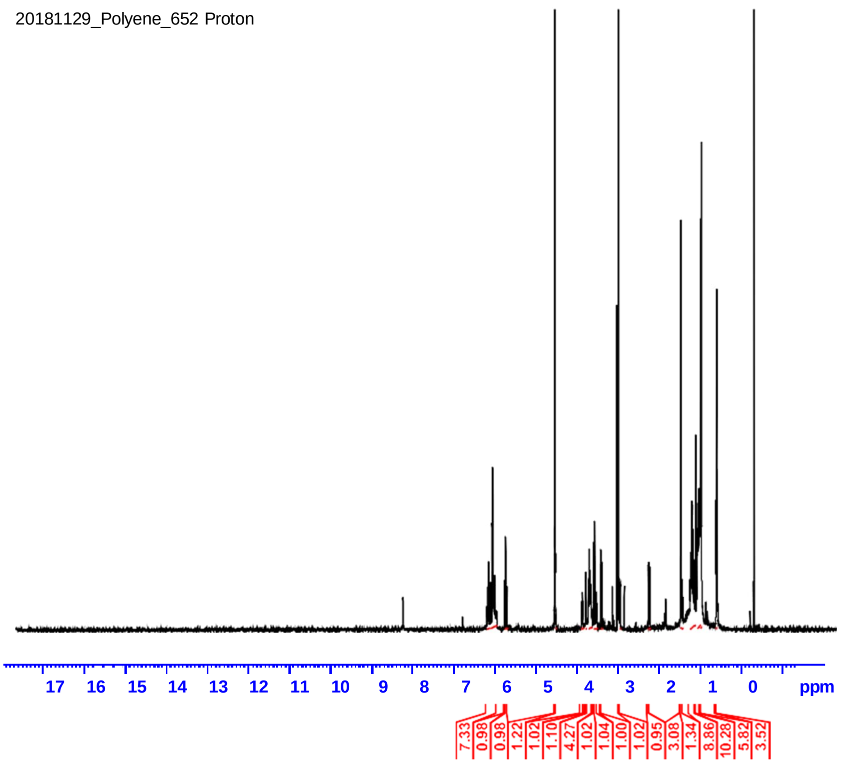


**Figure 2.** $^{1}$H spectrum of **1** in MEOD.

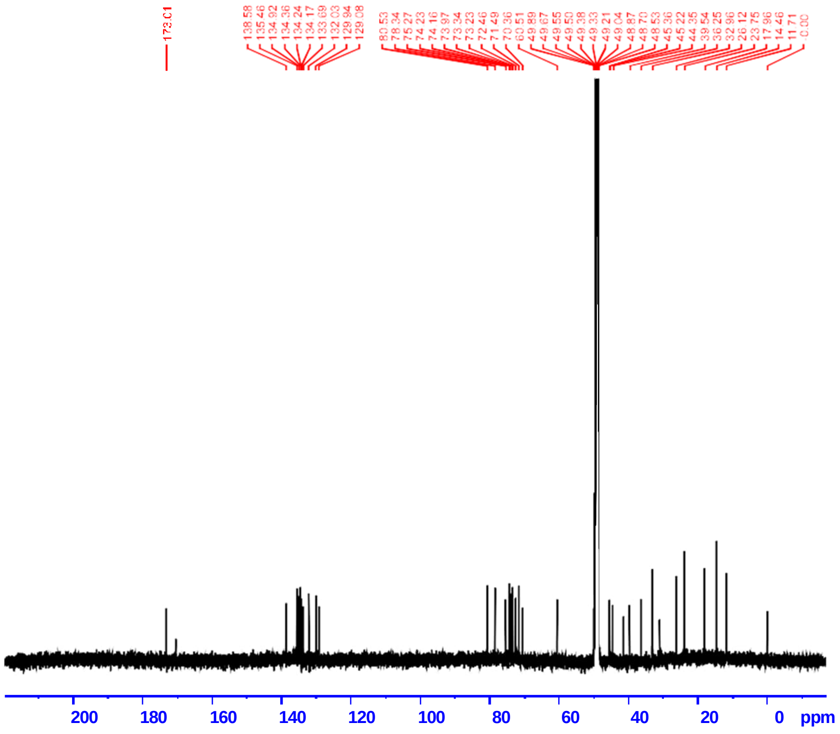


**Figure 3.** $^{13}$C spectrum of **1** in MeOD.

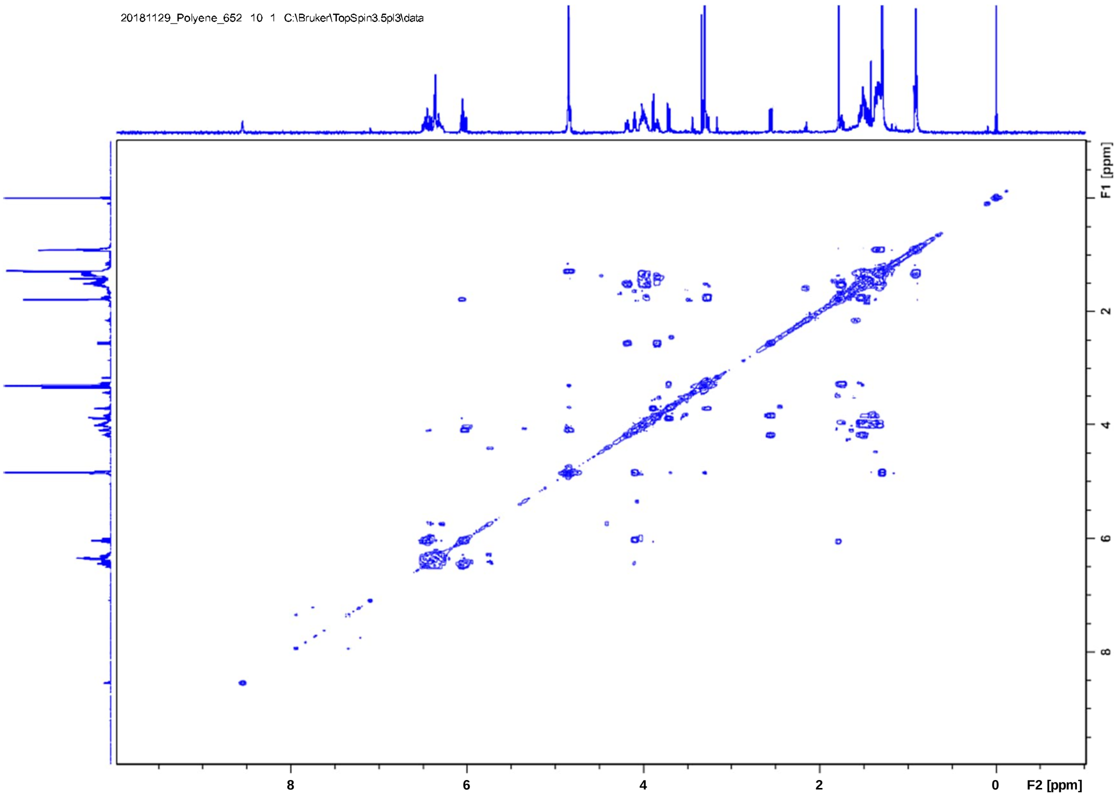


**Figure 4.** COSY spectrum of **1** in MEOD.

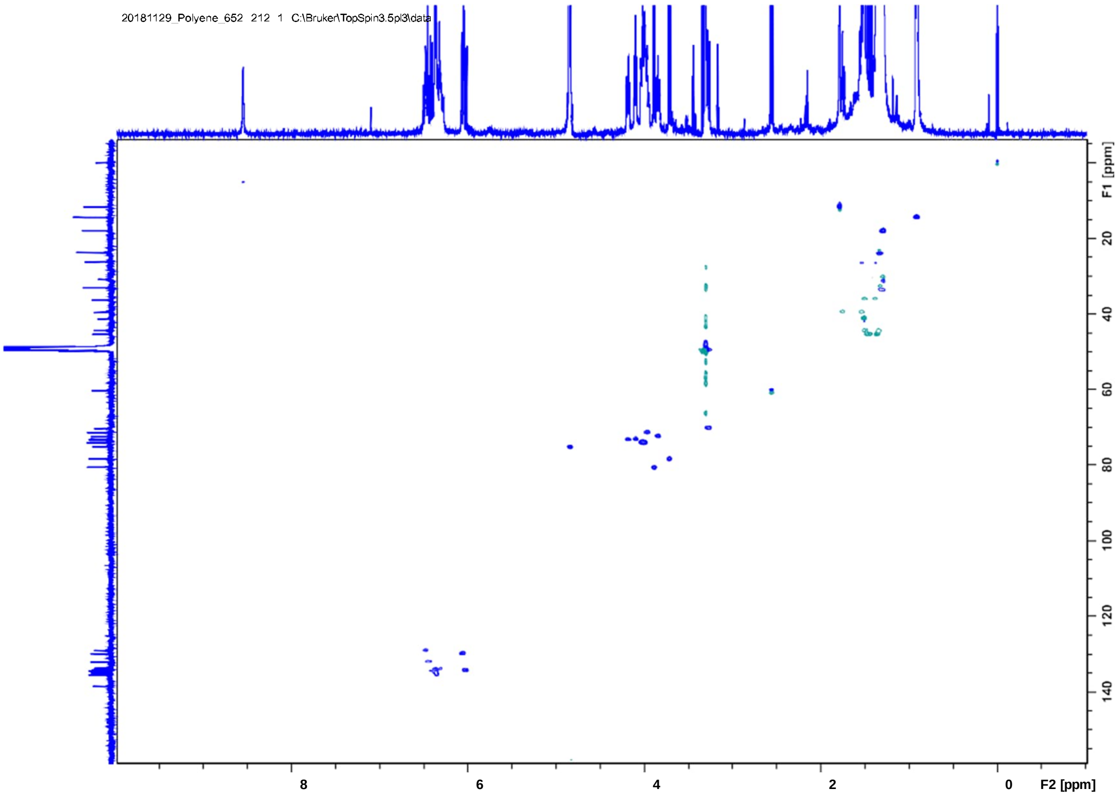


**Figure 5.** HSQCDE spectrum of **1** in MeOD.

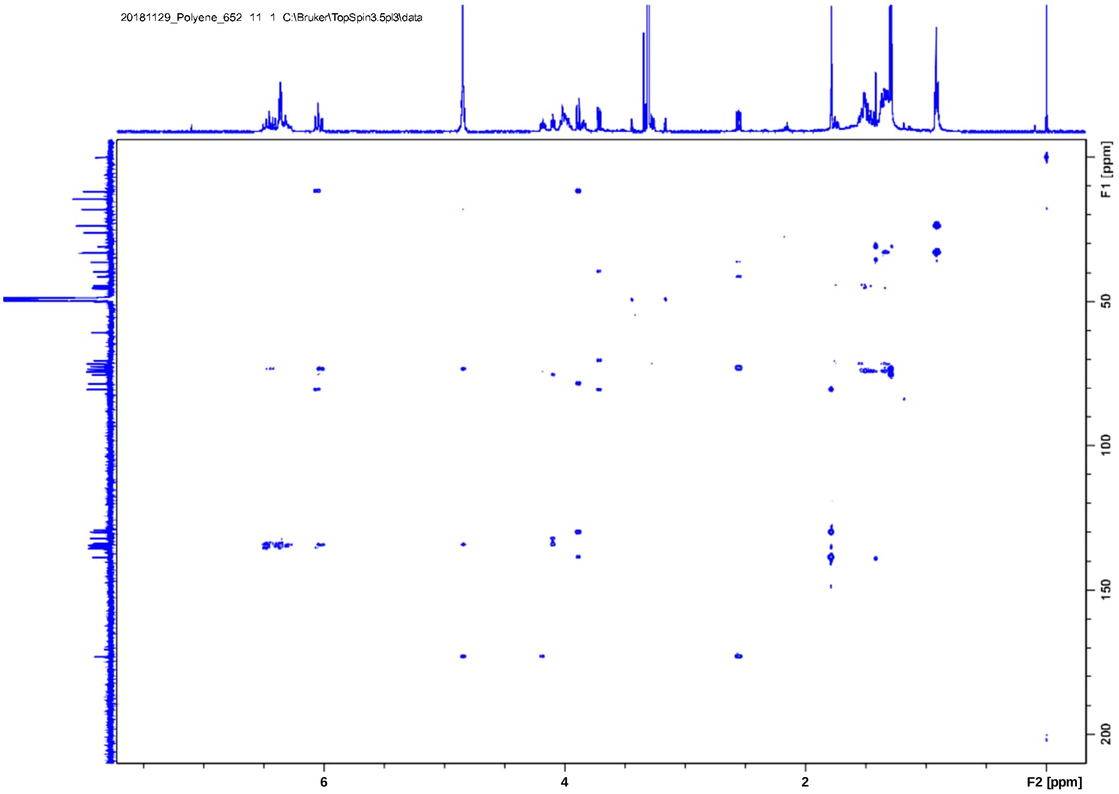


Figure 6. HMBC spectrum of **1** in MEOD.

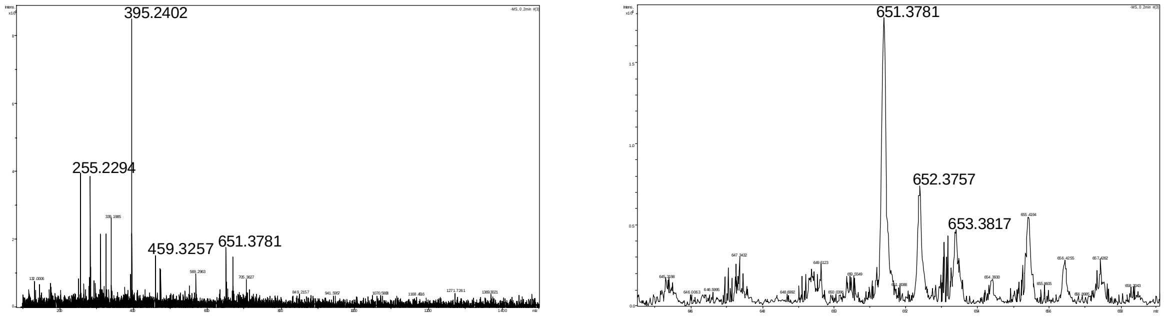


**Figure 7.** HR-MS spectrum of **1**. [H-M-H]$^-$ calc. 651.3783 obs. 651.3781.

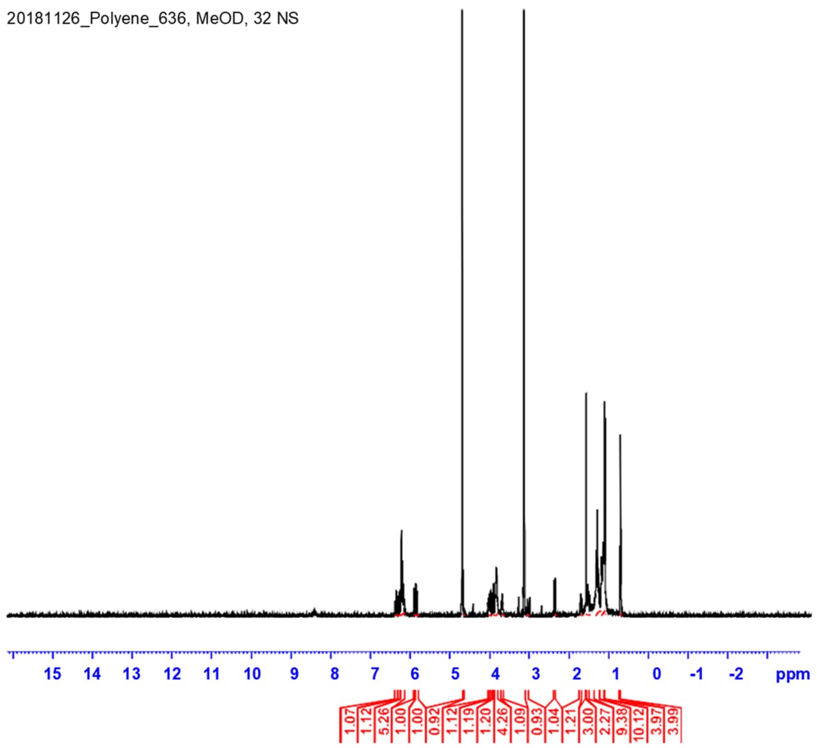


**Figure 8.** $^{1}$H spectrum of **2** in MEOD.

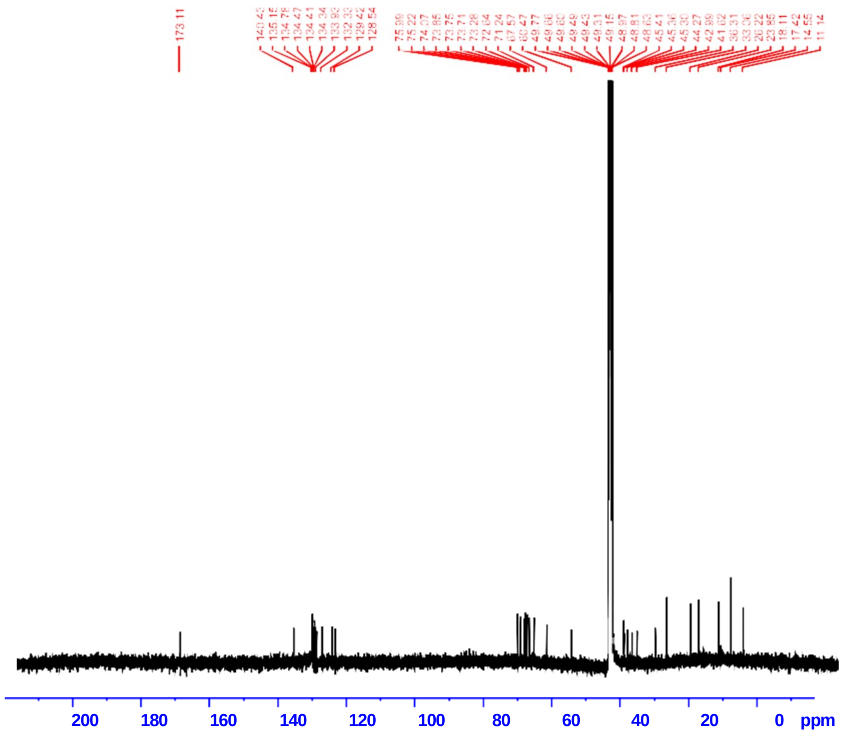


**Figure 9.** $^{13}$C spectrum of **2** in MeOD.

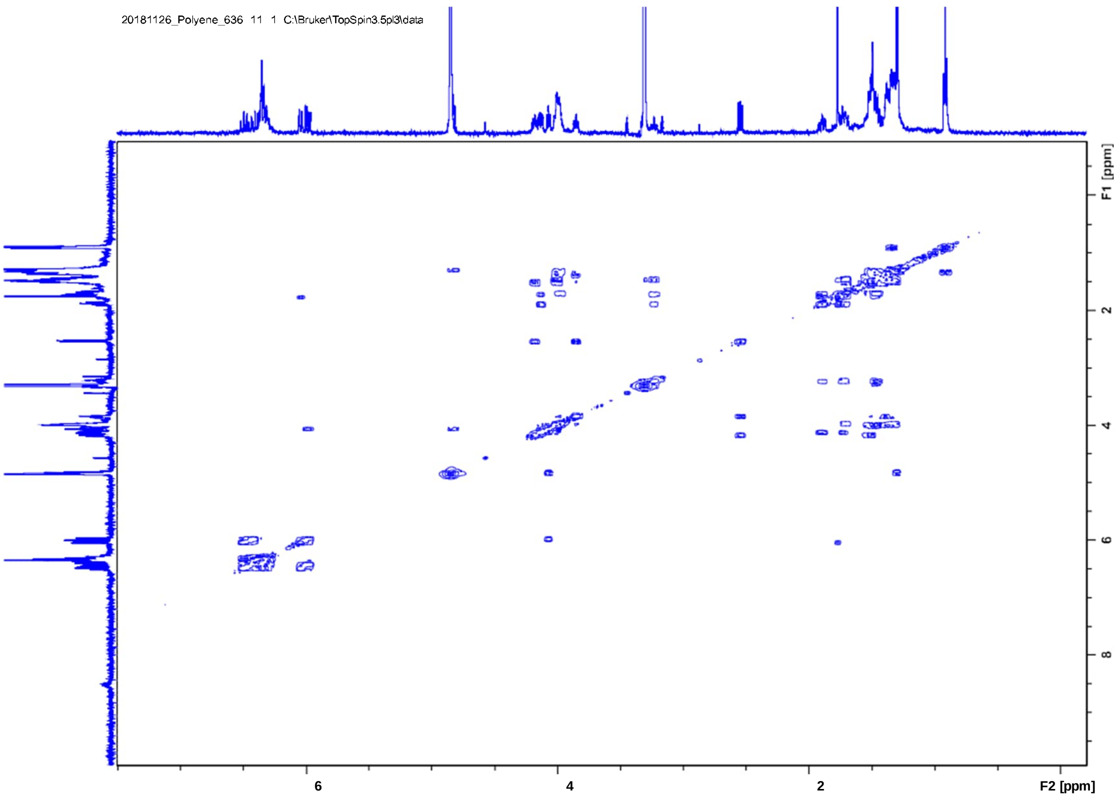


**Figure 10.** COSY spectrum of **2** in MeOD.

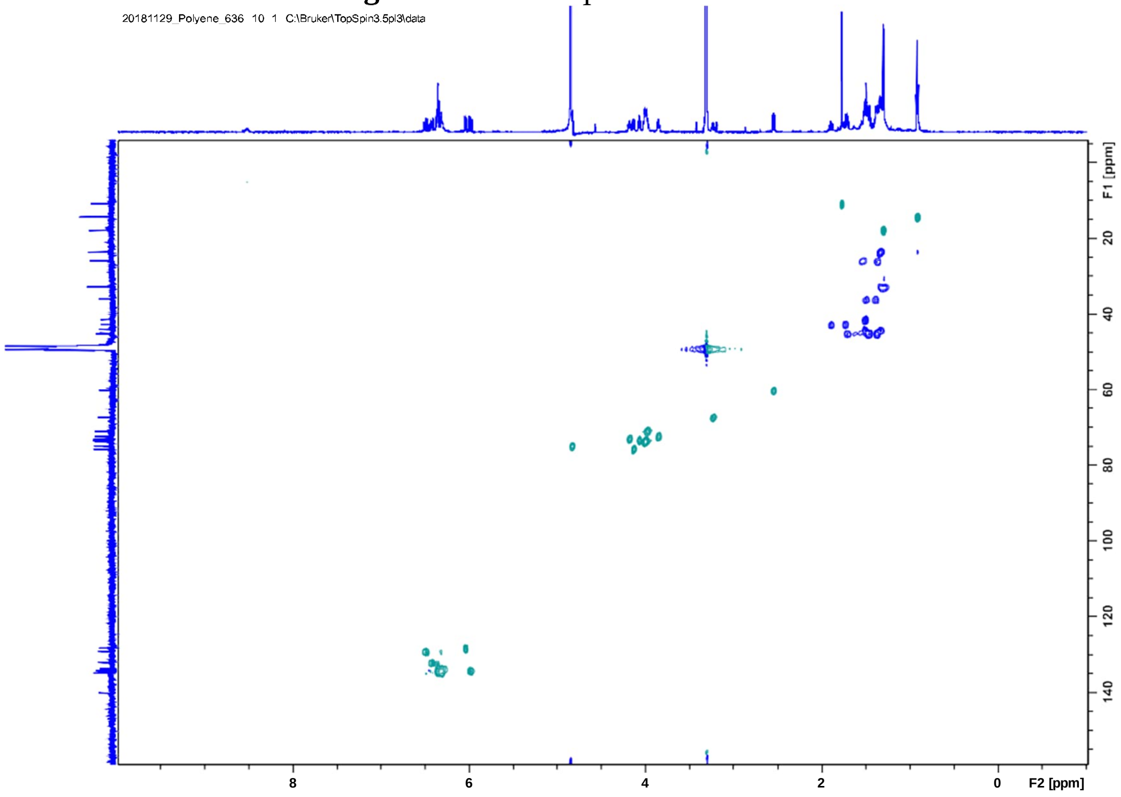


**Figure 11.** HSQCDE spectrum of **2** in MEOD.

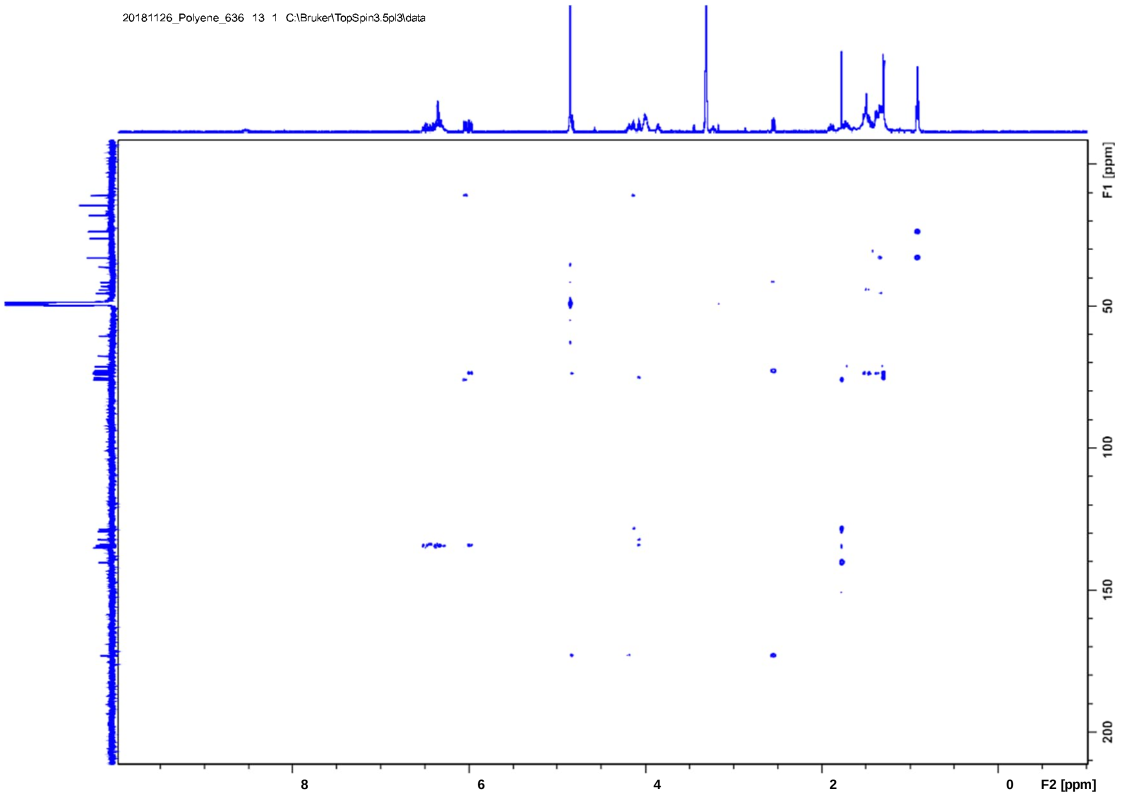


**Figure 12.** HMBC spectrum of **2** in MeOD.

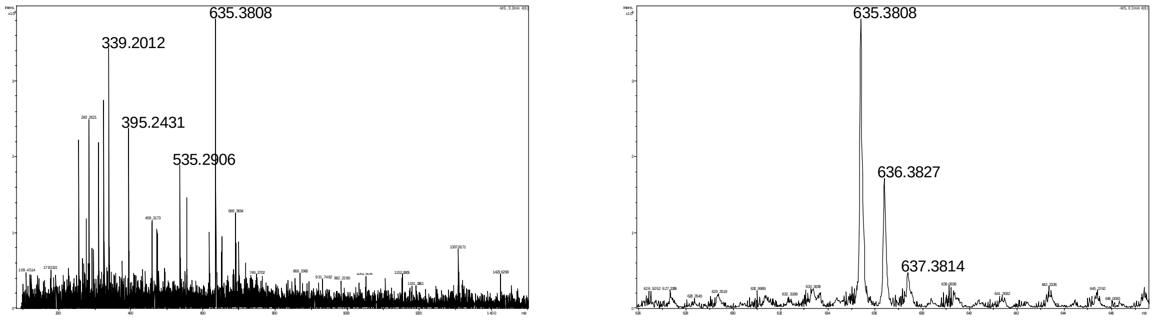


**Figure 13.** HR-MS spectrum of **2**. [H-M-H]$^{-}$ calc. 635.2958, obs. 635.3808.

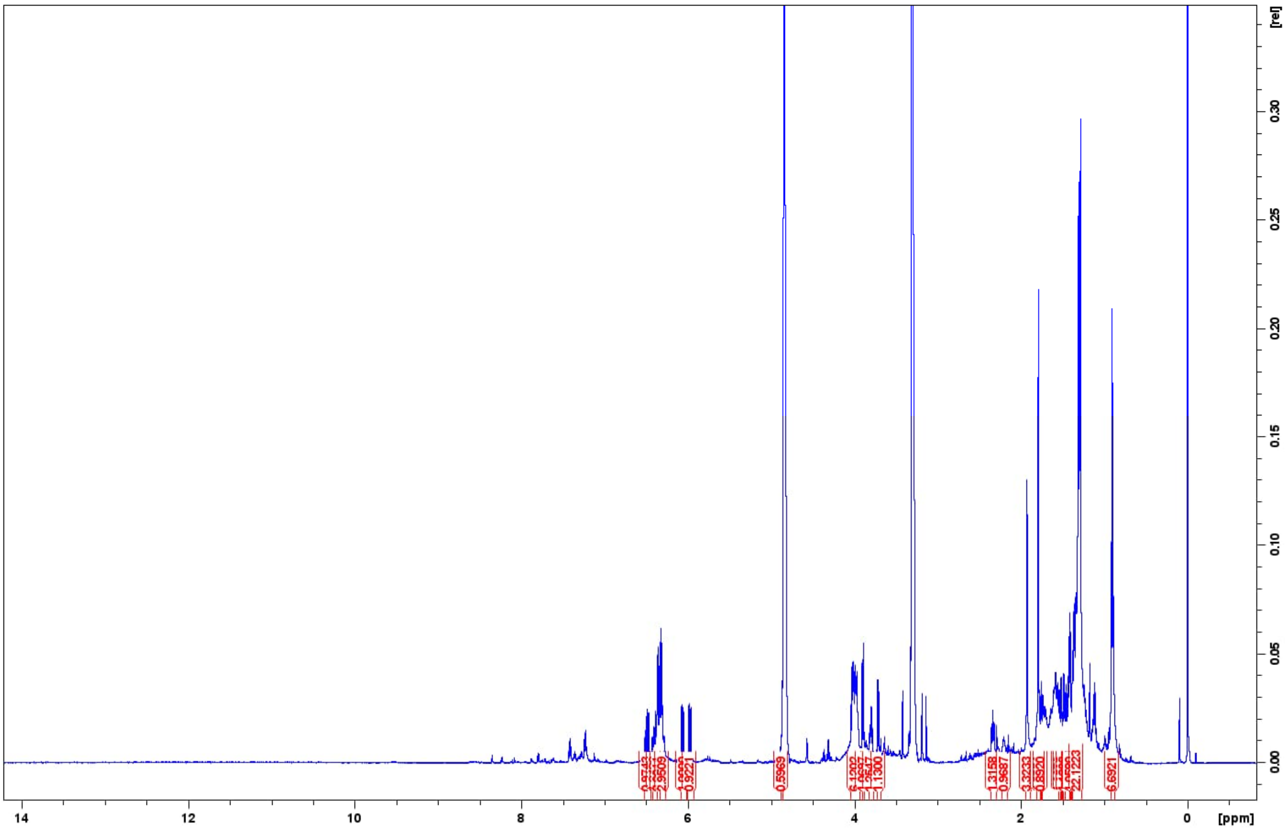


**Figure 14.** $^{1}$H NMR spectrum ($CD_3OD$, 600 MHz) of **3**.

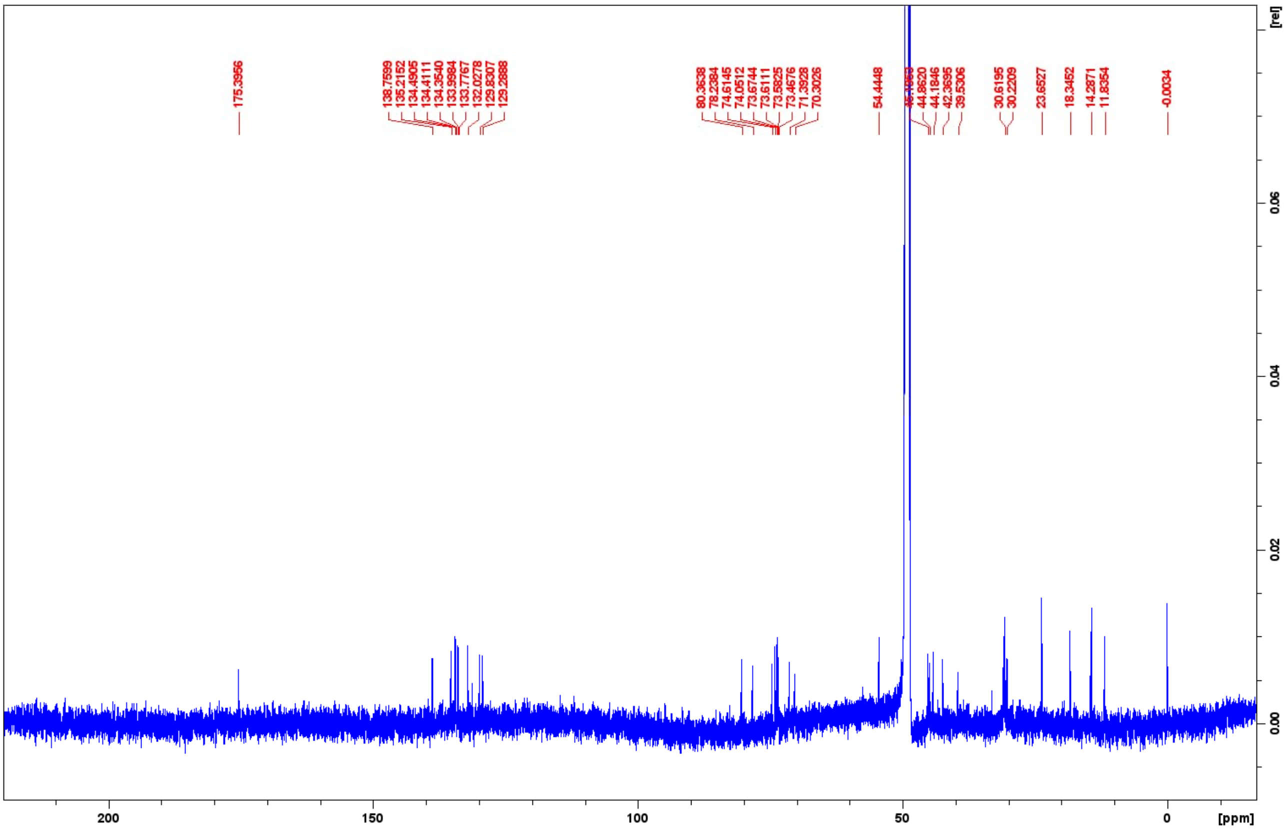


**Figure 15.** $^{13}$C NMR spectrum ($CD_3OD$, 125 MHz) of **3**.

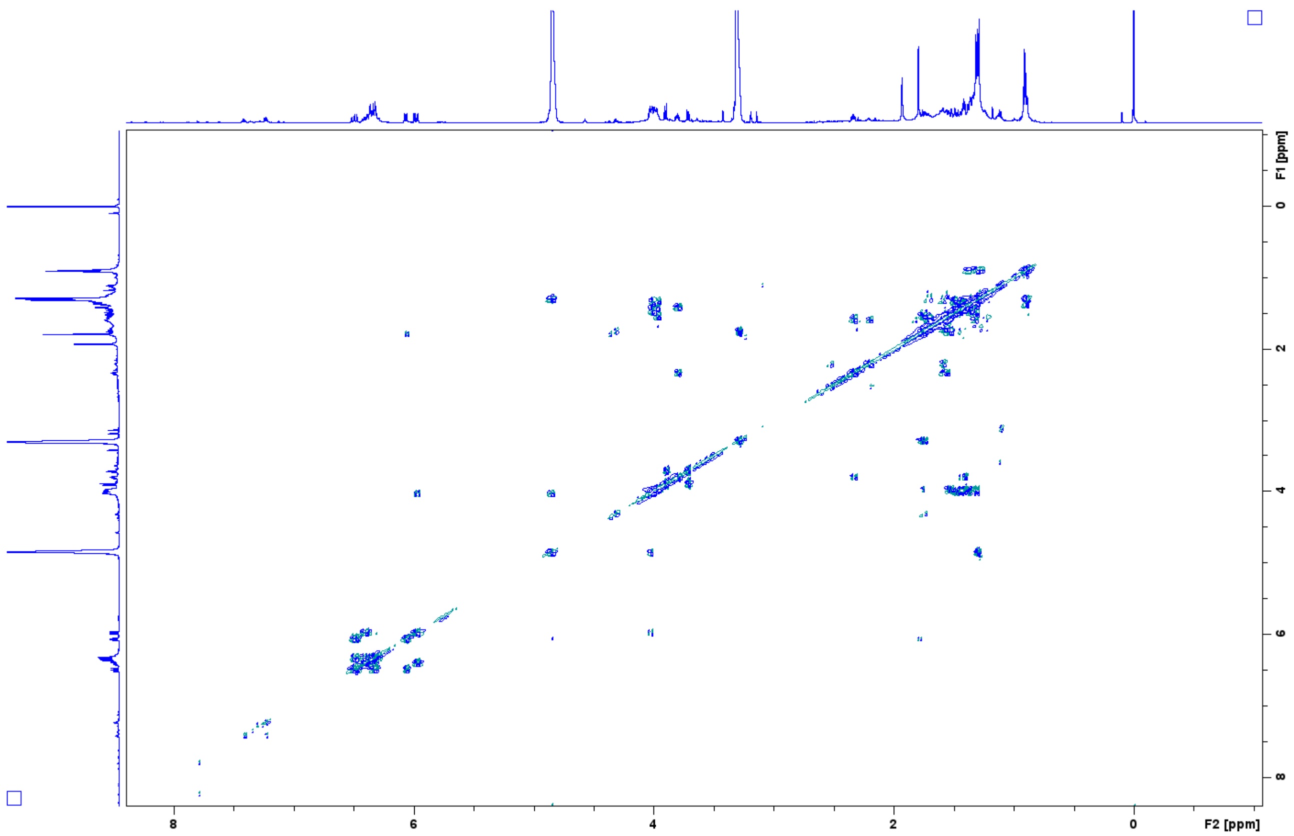


**Figure 16.** $^{1}H$, $^{1}H$-COSY spectrum ($CD_3OD$, 600 MHz) of **3**.

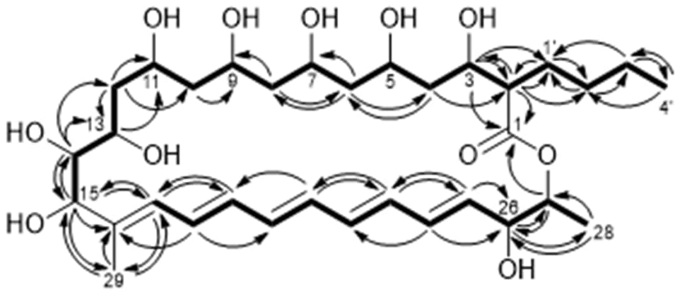


14-hydroxyisochainin

**Figure 17.** $^{1}H$, $^{1}H$-COSY (▬) and selected HMBC (→) correlations of **3**.

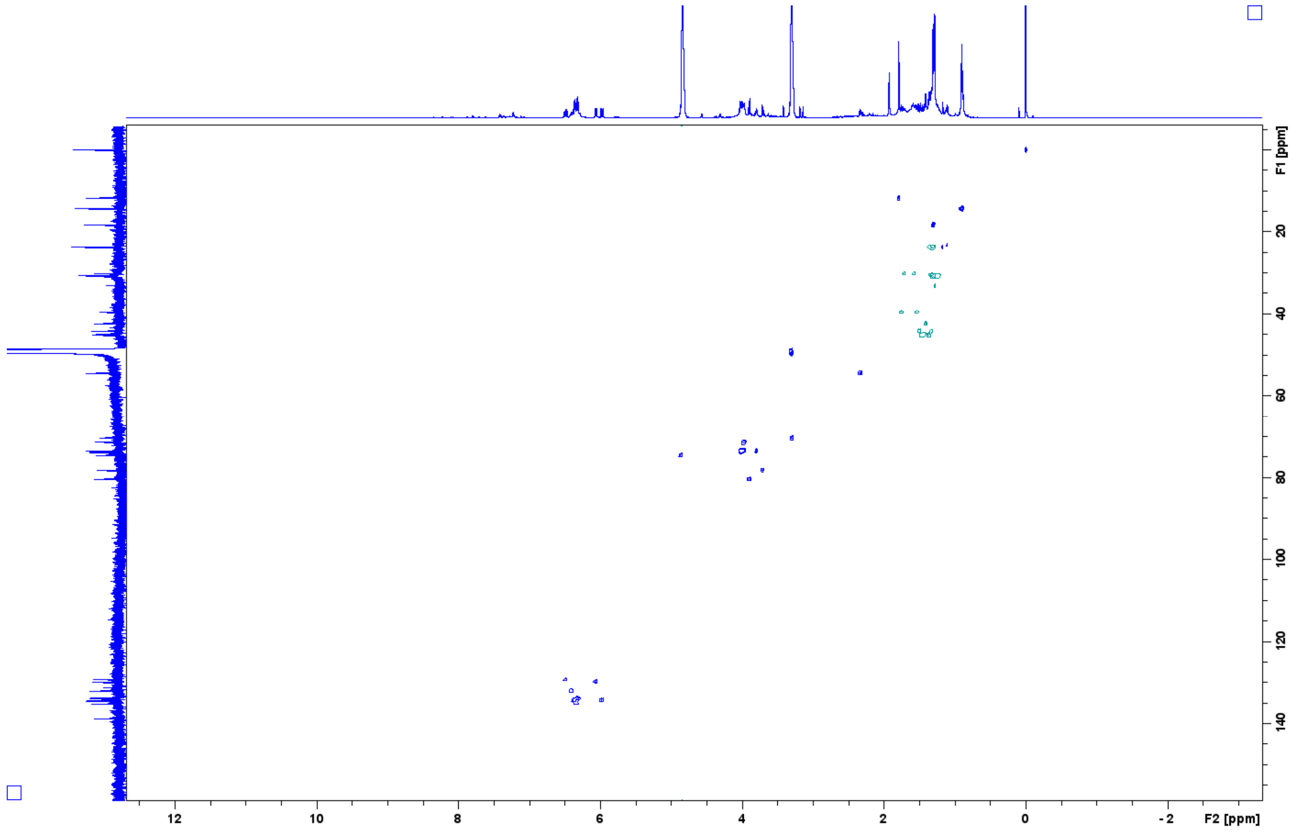


**Figure 18.** HSQC spectrum ($CD_3OD$, 600 MHz) of **3**.

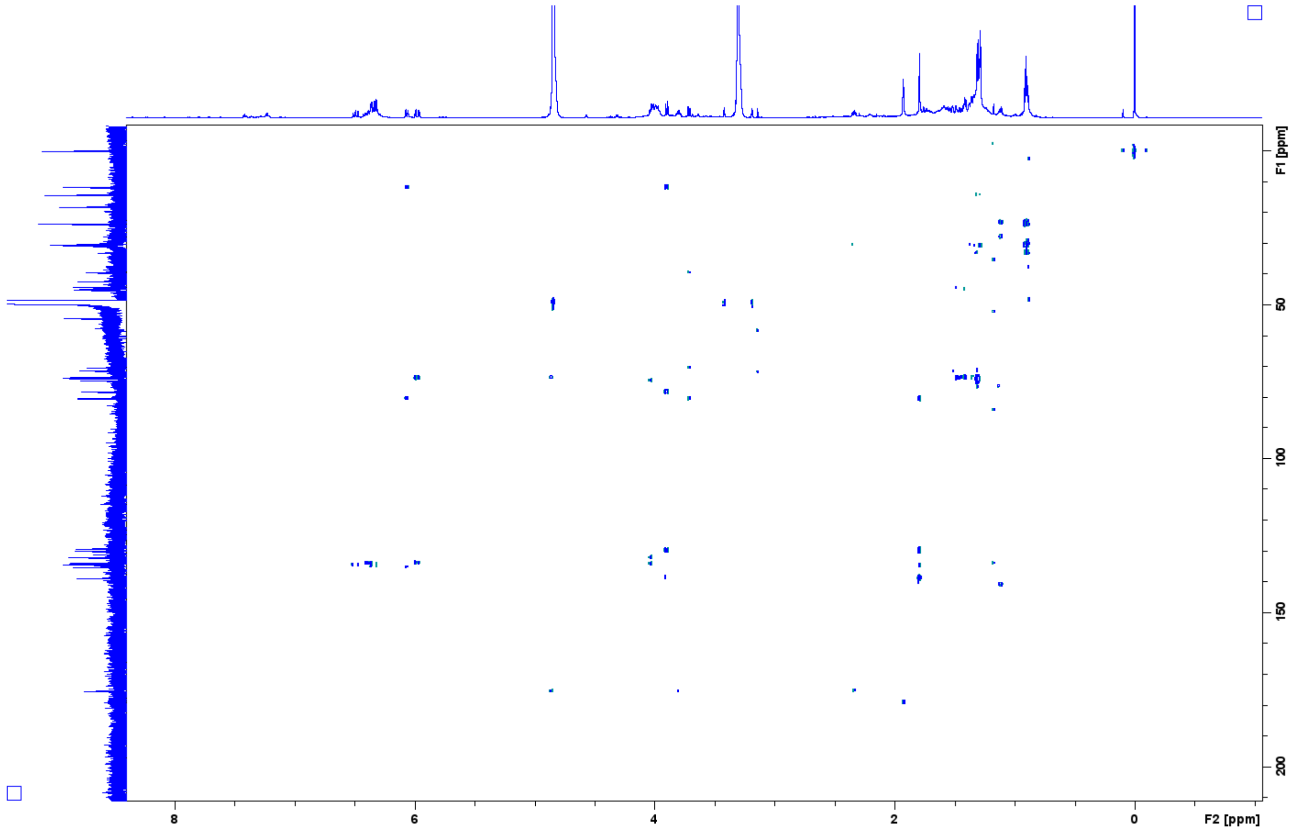


**Figure 19.** HMBC spectrum ($CD_3OD$, 600 MHz) of **3**.

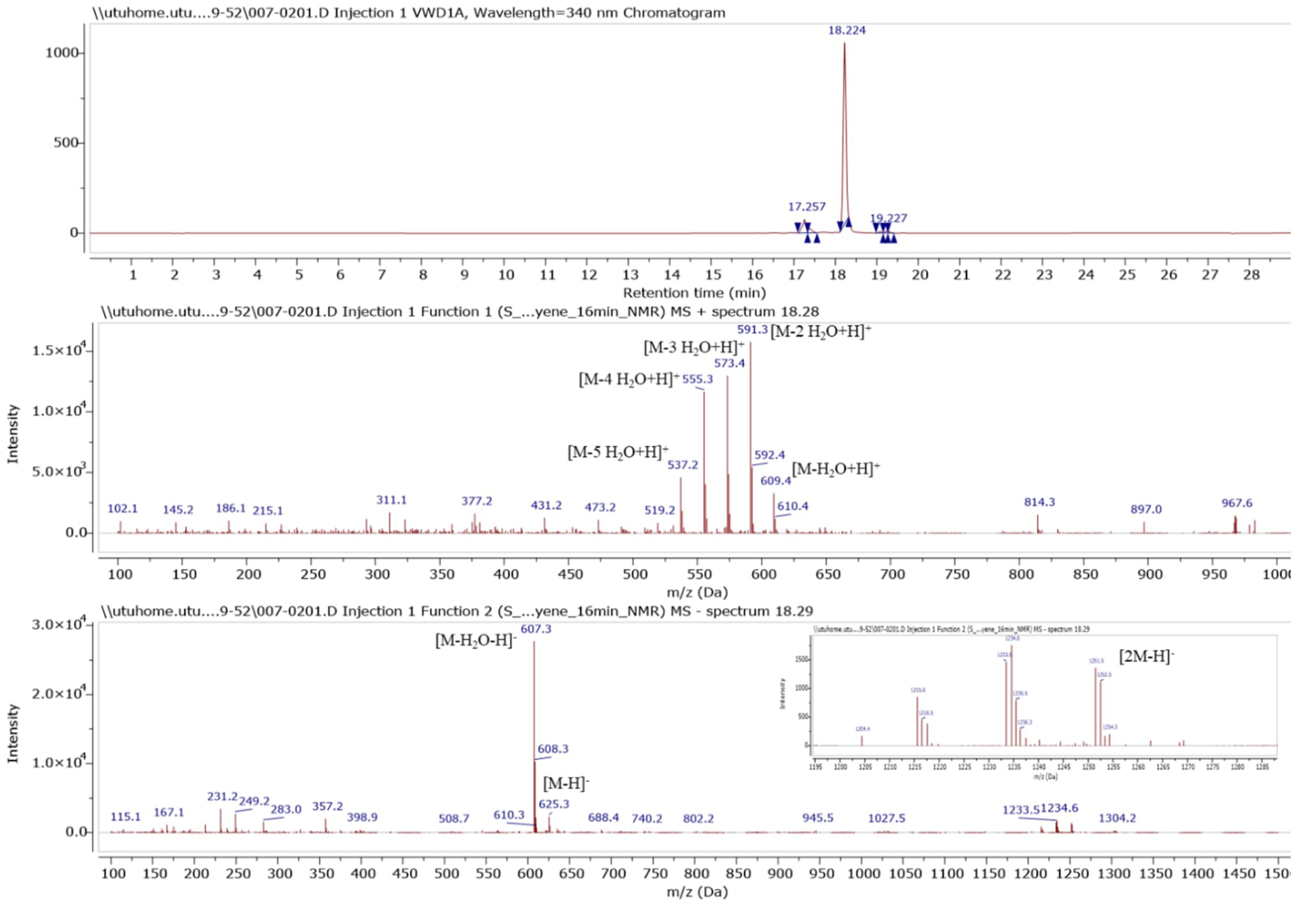


**Figure 20.** HPLC-MS analysis of **3**.

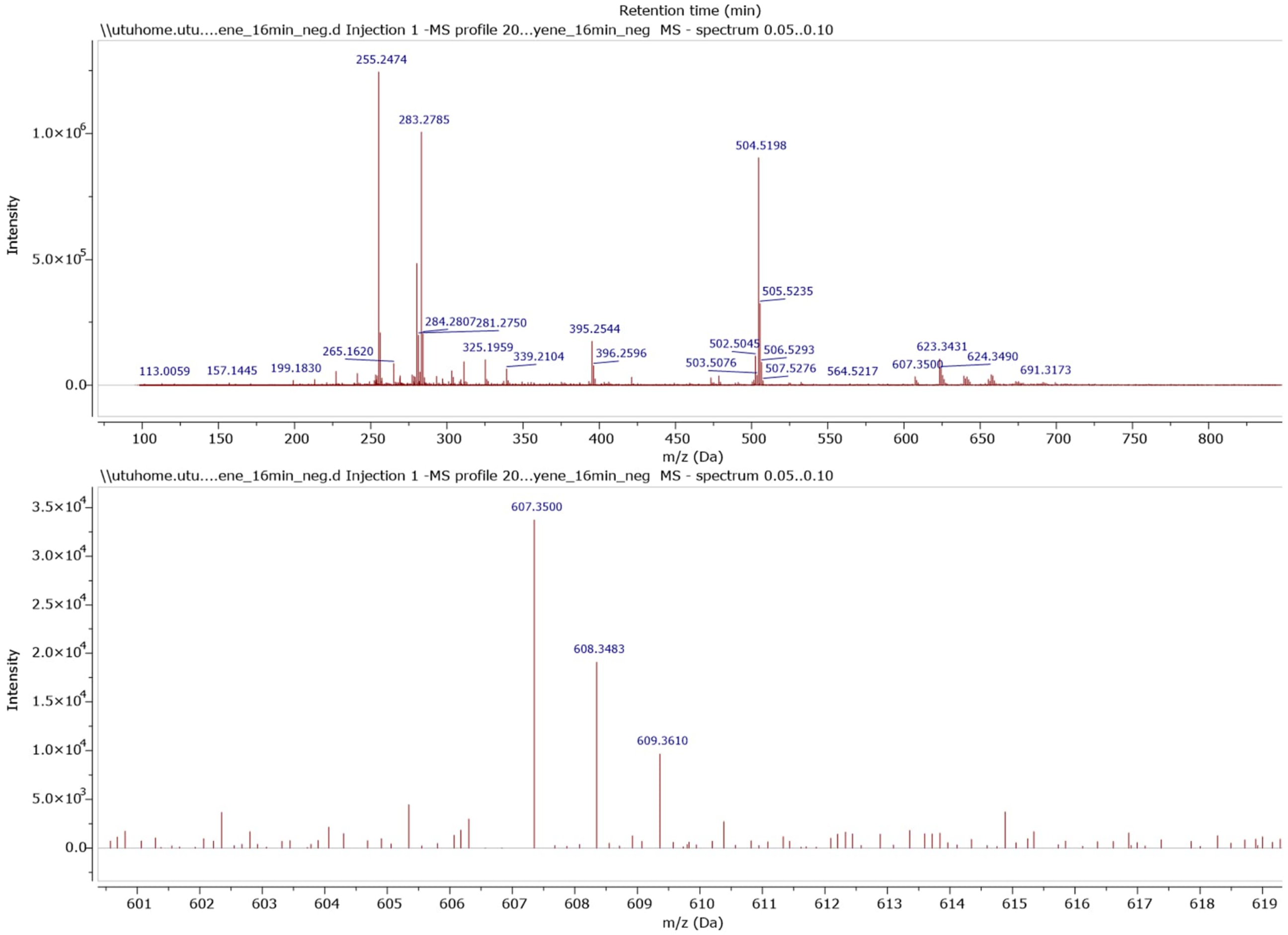


**Figure 21.** HR-MS spectrum of **3**. $[M-H_2O-H]^-$ calc. 607.3482, obs. 607.3500.

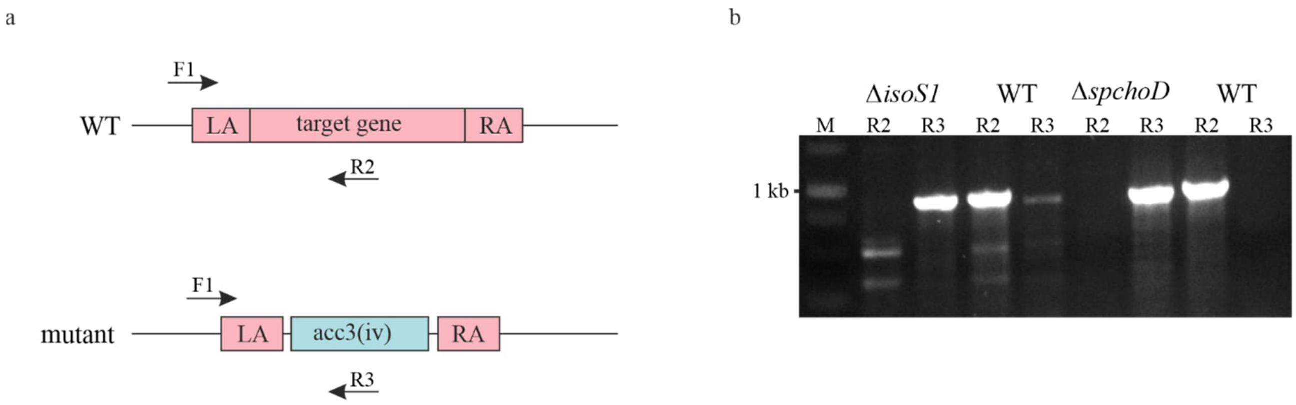


**Figure 22.** Confirmation of gene knockout by PCR analysis. **a,** Schematic representation of gene disruption in *S. peucetius* ATCC 27952 and the position of PCR primers. **b,** PCR amplification with two primer sets: the knockout-specific primers F1 and R3 resulted in the amplification of a single ≈ 900 bp fragment in both *ΔisoS1* and *ΔspchoD* mutants, while the use of wild-type-specific primers F1 and R2 in mutants did not produce a PCR product, indicating successful gene inactivation.
